# Multigenerational downregulation of insulin/IGF-1 signalling in adulthood improves lineage survival, reproduction, and fitness in *C. elegans* supporting the developmental theory of ageing

**DOI:** 10.1101/2020.08.19.257253

**Authors:** Elizabeth ML Duxbury, Hanne Carlsson, Kris Sales, Zahida Sultanova, Simone Immler, Tracey Chapman, Alexei A Maklakov

## Abstract

Adulthood-only downregulation of insulin/IGF-1 signalling (IIS), an evolutionarily conserved pathway regulating resource allocation between somatic maintenance and reproduction, increases lifespan without fecundity cost in the nematode, *Caenorhabditis elegans*. However, long-term multigenerational effects of reduced IIS remain unexplored and are proposed to carry costs for offspring quality. To test this hypothesis, we ran a mutation accumulation (MA) experiment and downregulated IIS in half of the 400 MA lines by silencing *daf-2* gene expression using RNA interference (RNAi) across 40 generations. Contrary to the prediction, adulthood-only *daf-2* RNAi reduced extinction of MA lines both under UV-induced and spontaneous mutation accumulation. Fitness of the surviving UV-induced MA lines was higher under *daf-2* RNAi. Reduced IIS increased intergenerational F1 offspring fitness under UV stress but had no quantifiable transgenerational effects. Functional *hrde-1* was required for the benefits of multigenerational *daf-2* RNAi. Overall, we found net benefit to fitness from multigenerational reduction of IIS and the benefits became more apparent under stress. Because reduced *daf-2* expression during development carries fitness costs, we suggest that our findings are best explained by the developmental theory of ageing, which maintains that the decline in the force of selection with age results in poorly regulated gene expression in adulthood.

## Introduction

Ageing, the physiological deterioration of an organism leading to increased probability of death and decreased reproduction with advancing adult age, is taxonomically ubiquitous but remains incompletely understood^1–4^. While there is broad agreement that ageing evolves because natural selection gradients on traits decline after reproductive maturity^5–9^, the proximate causes of late-life deterioration are less clear^1,2,10,11^. The “disposable soma” theory of ageing suggests that ageing evolves due to competitive resource allocation between the germline and the soma^12–14^ It follows that increased investment in somatic maintenance leading to longer lifespan trades-off with traits associated with reproduction, such as the number or size of progeny^15,16^. Despite considerable support for trade-offs between somatic maintenance and reproduction, growing empirical work questions the universality of such resource allocation trade-offs in the evolution of ageing^2,11,17,18^. Several studies have shown that experimentally increased lifespan, often via the downregulation of genes in nutrient-sensing signalling pathways in adulthood, is not detrimental to reproduction^2,18,19,20,21^. These results are in line with the hypothesis that a gradual decline in selection gradients with advancing age after reproductive maturity results in physiological processes that are optimised for successful early life reproduction, but start causing harm in later life, contributing to somatic deterioration^2,22–28^.

However, it has been proposed that reduced offspring quality, for example via reduced germline maintenance, can be a hidden cost of lifespan extension via reduced nutrient-sensing signalling^18^. Germline maintenance, the repair and surveillance of genomic and proteomic integrity in germline stem cells and gametes, is energetically expensive^18,29^ and germline signalling plays a key role in resource allocation to somatic maintenance^18,29,30,31^. Germline ablation results in increased somatic maintenance and lifespan in *Drosophila melanogaster* fruitflies^32^ and *Caenorhabditis elegans* nematodes^33–37^, although lifespan extension in *C. elegans* requires an intact somatic gonad^33^. It is important to note that the requirement of the intact somatic gonad for lifespan extension in germline-less worms does not negate the possibility of a resource allocation trade-off as sometimes implied, but only shows that germline signalling is required to mediate the effect^18,32,34^. Similarly, recent work in *Danio rerio* zebrafish suggests that germline ablation increases somatic maintenance under stress^38^. Furthermore, nutritional stress in *C. elegans* results in germline reduction and increased lifespan^39^. When soma-to-germline communication is disrupted, the number of germ cells is unaffected and lifespan extension is abolished^39^. Taken together, these results suggest that germline maintenance is costly and can trade-off with life-history traits such as lifespan and offspring number^18,29^. The corollary is that increased investment in somatic maintenance and longevity can trade-off with offspring quality by increasing germline mutation load in offspring.

We tested this hypothesis by focusing on lifespan extension via reduced insulin/IGF-1 signalling (IIS) in *Caenorhabditis elegans*. IIS is an evolutionarily conserved pathway that regulates the physiological response of organisms to their environment^3^. IIS mechanistically links nutrient intake with development, growth, reproduction and lifespan across diverse taxa^3,31,40^. Reduced IIS, via genetic and environmental interventions, consistently extends lifespan^31^. Mutations that reduce the function of *daf-2* gene in *C. elegans,* the homologue of human insulin/IGF-1 receptor, increase lifespan and reduce fitness^41^. However, previous work showed that reducing IIS in adulthood via *daf-2* RNA interference (RNAi) knockdown in a single generation increases lifespan without a cost to offspring number^19^, quality^20^ or fitness in variable environments^28^. It is possible, however, that multi-generational *daf-2* knockdown will expose a hidden cost to offspring number and/or offspring quality via accumulation of germline mutations. Here we tested this prediction via adulthood-only *daf-2* RNAi knockdown for 40 consecutive generations in 400 spontaneous and UV-induced mutation accumulation (MA) lines. We found that multi-generational adulthood-only *daf-2* knockdown reduced lineage extinction under both spontaneous and UV-induced MA and increased fitness of remaining lineages under UV-induced MA. Reduced IIS protected MA lines from infertility, egg hatching failure and developmental abnormalities and increased offspring fitness under UV irradiation, but we found no evidence of trans-generational effects of adulthood-only *daf-2* RNAi.

## Materials and Methods

Supplementary methods details are included in the Supplementary Material.

### Nematode stocks and culture

The nematode (roundworm) *Caenorhabditis elegans* N2 wild-type (Bristol) and heritable RNAi deficiency 1 (*hrde-1*) mutant strains were defrosted from stocks acquired from Caenorhabditis Genetics Center (University of Minnesota, USA, funded by NIH Office of Research Infrastructure Programs, P40 OD010440) and from the lab of Prof. Eric Miska (Gurdon Institute, University of Cambridge, UK), respectively, and stored at −80°C until use.

Defrosted *C. elegans* strains were reared through two generations prior to setup, on NGM agar supplemented with the fungicide nystatin and antibiotics streptomycin and ampicillin, to prevent infection (each at 100 μg/ml, as standard recipes, e.g.^42^) and seeded with the antibiotic-resistant *Escherichia coli* bacterial strain OP50-1 (pUC4K, from J. Ewbank at the Centre d’Immunologie de Marseille-Luminy, France). We bleached eggs from the grandparents of experimental individuals, to standardise parental age and remove any infection or temperature effects from defrosting, prior to experiments.

### Reducing IIS via *daf-2* RNAi feeding in adulthood

To downregulate adulthood expression of the insulin-like sensing signalling receptor homolog, *daf-2*, we fed late-L4 larvae with *Escherichia coli* bacteria expressing *daf-2* double-stranded RNA, that decreases mRNA levels of the complementary transcribed *daf-2* systemically^43,44^. The *daf-2* gene is upstream of and inhibits the action of master regulator daf-16/FOXO. RNase-III deficient, IPTG-inducible HT115 *E. coli* bacteria with an empty plasmid vector (L4440) was used as the control (as^19,20,44^). RNAi clones were acquired from the Vidal feeding library (Source BioScience, created by M. Vidal lab, Harvard Medical School, USA) and all clones were verified via sequencing prior to delivery.

### Ultraviolet wavelength C (UV-C) irradiation

To induce mutations and stress, we UV-irradiated Day 2 adults, with a calibrated ultraviolet-C (UV-C) radiation dose of 46J/m^2^ (wavelength 254nm), via 20 second exposure to the UV-C radiation emitted from the lamp of a Thermo Scientific Heraguard ECO Safety Cabinet (calibration details in Supplementary Methods). This dose is in the range of previous UV-C irradiation doses for *C. elegans* adults or eggs^45–48^. Our pilot work showed that this dose reduced the fecundity of Day 2 adults laying at 20°C by 61% compared with unexposed sham controls (n=30 worms/treatment, mean Day 2 offspring count = 35 for UV, = 90 for non-UV; full data not shown). Non-irradiated control worms received a sham-irradiation, by being positioned in identical orientation under the UV-C lamp for 20 seconds, while it was switched off.

### Mutation accumulation (MA) lines

To determine the effects of reduced adulthood IIS, via *daf-2* RNAi, on spontaneous and UV-induced mutation accumulation, we established 800 MA lines, across two genetic backgrounds. The eight experimental treatments were the full-factorial combinations of genotype (N2 or *hrde-1* mutant), RNAi treatment (empty vector or *daf-2* RNAi) and irradiation (UV or sham), with 100 MA lines per treatment. We ran each of the eight treatments in parallel, divided into two time-staggered independent blocks of 50 lines per treatment, for logistical reasons and to capture any between-block variation. We tracked MA lines until the majority of UV-irradiated lines were extinct-40 generations for N2 and 25 generations for *hrde-1* mutants. Every MA generation, *daf-2* RNAi or empty vector was delivered from the start of adulthood onwards, and all lines developed on empty vector. More details are provided in the Supplementary Methods.

Each MA line was propagated as a single individual hermaphrodite per generation, to create successive genetic bottlenecks, allowing *de novo* deleterious mutations to accumulate in the relative absence of selection^49–51^, if they had no effect on developmental time or viability, and in the absence of mating. MA is a common approach used in several species, to study the evolutionary genetics of *de novo* mutation rates^52^. In wild type *C. elegans,* the spontaneous mutation rate in MA lines is positively correlated with the rate of fitness decay^53^.

### Validation of *daf-2* gene expression under RNAi

We quantified, via quantitative reverse transcriptase polymerase chain reaction (qRT-PCR), the extent of downregulation of *daf-2* expression in Day 2 adults from the MA lines at generation 20, following adulthood-only *daf-2* RNAi knockdown, relative to generation 20 empty vector control MA lines. A sample from all eight MA treatments above that had been frozen at generation 20 were used (mean n= 6 MA lines per treatment, with mean n=3 individual worms per MA line; Table S1). Previous work has shown that adulthood *daf-2* RNAi reduces *daf-2* expression in Day 2 N2 adults by 50%, relative to controls^28^, equivalent to the first generation of MA. The aim was to test the degree of *daf-2* knockdown after 20 generations of *daf-2* RNAi and MA in both N2 and *hrde-1* mutant backgrounds. Worms for the gene expression assay were maintained under standard conditions (as the main experiment) and were reared for two generations on empty vector prior to the assay, to remove freezing effects and age-synchronise. RNAi was applied from late-L4 stage. Further method details are included in Supplementary Material.

### Effect of 20 generations of spontaneous and UV-induced mutation accumulation on fitness and life history of surviving MA lines

To test for life history differences between the N2 wild-type MA lines after 20 generations of MA and of adulthood *daf-2* RNAi, we assayed age-specific reproduction, egg size, male production and adult heat shock resistance (n=100 grand-offspring from each of the four N2 MA treatments). Further details in Supplementary Methods.

Individual age-synchronised, late-L4 hermaphrodites were set up on separate e.v. plates, labelled with a unique identifier, to blind experimenters to treatment identity, and thus avoid bias. The two experimenters conducted the life history assay in two simultaneous time blocks and each block had an identical and equal representation of individual worms from all four treatments (n=50 per treatment per block).

For all experiments, we assayed age-specific offspring production (fecundity) over the first 6 days of adult life (the reproductive window for *C. elegans* hermaphrodites), by transferring to fresh plates every 24h, to acquire daily reproduction counts, following the standard temporal resolution for *C. elegans* studies^19,20,54^. Plates were scored for viable adult offspring 2.5 days later. Individual fitness (lambda) was calculated- a measure analogous to the intrinsic rate of population growth, that is weighted by early reproduction and fast development ^15,20,55,56^. Total lifetime reproduction (lifetime reproductive success) was calculated for each individual as the sum of age-specific offspring counts across the first 6 days of adulthood.

To determine if reduced IIS influenced parental resource allocation into eggs under MA, we measured egg area (in mm^2^), with 75 samples per treatment. Eggs measured were laid by the individuals in the generation 20 age-specific reproduction assay at peak reproduction (Day 2). Egg size is commonly used as a proxy for parental investment^20,57^. We photographed one egg laid per individual, within 2 hours of lay, under 12x magnification light microscopy (Leica M165C with Lumenera Infinity 2-7C digital microscope camera).

### Transgenerational effects of *daf-2* RNAi on offspring fitness in N2 and *hrde-1* mutant backgrounds

To determine whether *daf-2* RNAi was transferred transgenerationally via the germline, we assayed, in a separate experiment, whether offspring fitness benefits persisted across three generations of offspring (F1, F2 and F3) from Day 2 parents treated with *daf-2* RNAi or empty vector control throughout adulthood, in N2 wild-type and *hrde-1* mutant backgrounds. Offspring of each generation were maintained on empty vector (n=30 individuals per treatment and per genotype) and assayed for daily reproduction.

### Effects of reduced IIS on parental fitness and total reproduction under intermittent fasting

To determine whether there was a reproductive cost of adulthood-only *daf-2* RNAi under resource limitation, we exposed N2 wild-type adults to either an intermittent fasting regime consisting of 9 hours of starvation on Days 1, 3, 5 and 7 of adulthood, or ad libitum feeding throughout life, and assayed daily reproduction. Full details in Supplementary Material.

### Statistical analyses

All analyses were conducted in R version 4.0.0^58^. Fitness and total lifetime reproduction data were plotted using the R package ‘dabestr’ (data analysis using bootstrap-coupled estimation^59^), for visualisation.

Parental survival in the intergenerational (first) experiment was analysed using a Cox proportional hazards mixed effects model^60^ (‘coxme’), fitting RNAi, UV and their interaction as fixed effects and plate as a random effect. Separate analyses were conducted with matricides either classed as deaths or as censors. The age at death response variable contained a coding variable to distinguish deaths from censored individuals (due to accidental losses). Extinction (survival trajectories) under mutation accumulation was analysed using Cox proportional hazards regression analysis (‘coxph’ function in ‘survival’ package). The generation number at which each MA line went extinct was the response variable (instead of age at death) and this contained a coding variable to distinguish extinctions from censors, as above. To test the effect of experimental block or genotype (either N2 wildtype or *hrde-1* mutant) on extinction and on the interaction between RNAi and UV effects under MA, maximal models were fitted with either a three-way RNAi x UV x block interaction, or a RNAi x UV x genotype interaction, and step-wise model simplification conducted^61^. Block was included as a fixed effect to test for repeatability between the two blocks. For all survival analyses, the z-scores and p-values were reported, with the z-score defined as the log of the hazard ratio (risk of death, exp(coef)) divided by the standard error of the log hazard ratio.

Age-specific reproduction was analysed using generalised linear mixed effects models to account for temporal pseudoreplication of repeated measures on the same individuals across lifetime, with the template model builder package (‘glmmTMB’ in R^62,63^). Models fitted with Poisson, Zero-Inflated Poisson, Generalised Poisson and Zero-Inflated Generalised Poisson error structure were compared using AIC values of model fit (‘AICtab’ function in ‘bbmle’ package), following^63^, to account for under- or over-dispersion and for zero-inflation, when it was found to occur in simulated residuals generated with the ‘DHARMa’ package^64^. Age and a quadratic form of age (age^2^) were fitted as fixed effects in both the conditional and zero-inflation model formula, and significance assessed in each case. The age^2^ term controlled for the curved (non-linear) trajectory of reproduction across age^65^. UV irradiation, RNAi treatment and their three-way interaction with age or age^2^ were fitted as fixed effects, and experimental block, observer and a unique plate identifier as random effects, the latter to control for repeated measures of fecundity from the same individuals across lifetime. Genotype was substituted with UV as a fixed effect to assess the transgenerational effects of *daf-2* RNAi on N2 versus *hrde-1* mutants. Models that did not converge were not included in model comparison, and the converging model with best AIC fit was presented (as^63^).

Fitness (lambda) was calculated using the ‘lambda’ function in the ‘popbio’ package. More details are provided in the Supplementary Material. Fitness, total reproduction and egg size data were analysed using a generalised linear model (GLM) with Gaussian error structure (‘lm’ function in ‘stats’ package). Locomotion, a measure of stress resistance to heat shock, was coded as a binary response variable (normal or abnormal movement)) and analysed using a GLM with binomial error structure.

## Results and Discussion

### Parental *daf-2* RNAi in adulthood extends lifespan and improves offspring fitness under UV-induced stress

Adulthood-only *daf-2* RNAi in N2 wildtype *C. elegans* nematodes, significantly extended parental lifespan relative to empty vector (e.v.) controls under benign conditions (no irradiation) and under ultraviolet-C (“UV”) irradiation-induced stress (Cox proportional hazards mixed effects model, coxme, with matricides censored, RNAi: z=10.530, p<0.001; UV: z=0.070, p=0.940; RNAi x UV: z= 1.500, p=0.130; matricides classed as dead: RNAi: z=10.400, p<0.001; UV: z=0.250, p=0.800; RNAi x UV: z= 1.310, p=0.190; Fig. 1a). This UV-induced stress encompassed both the mutagen properties of UV radiation and the stress response induced.

**Fig. 1.**
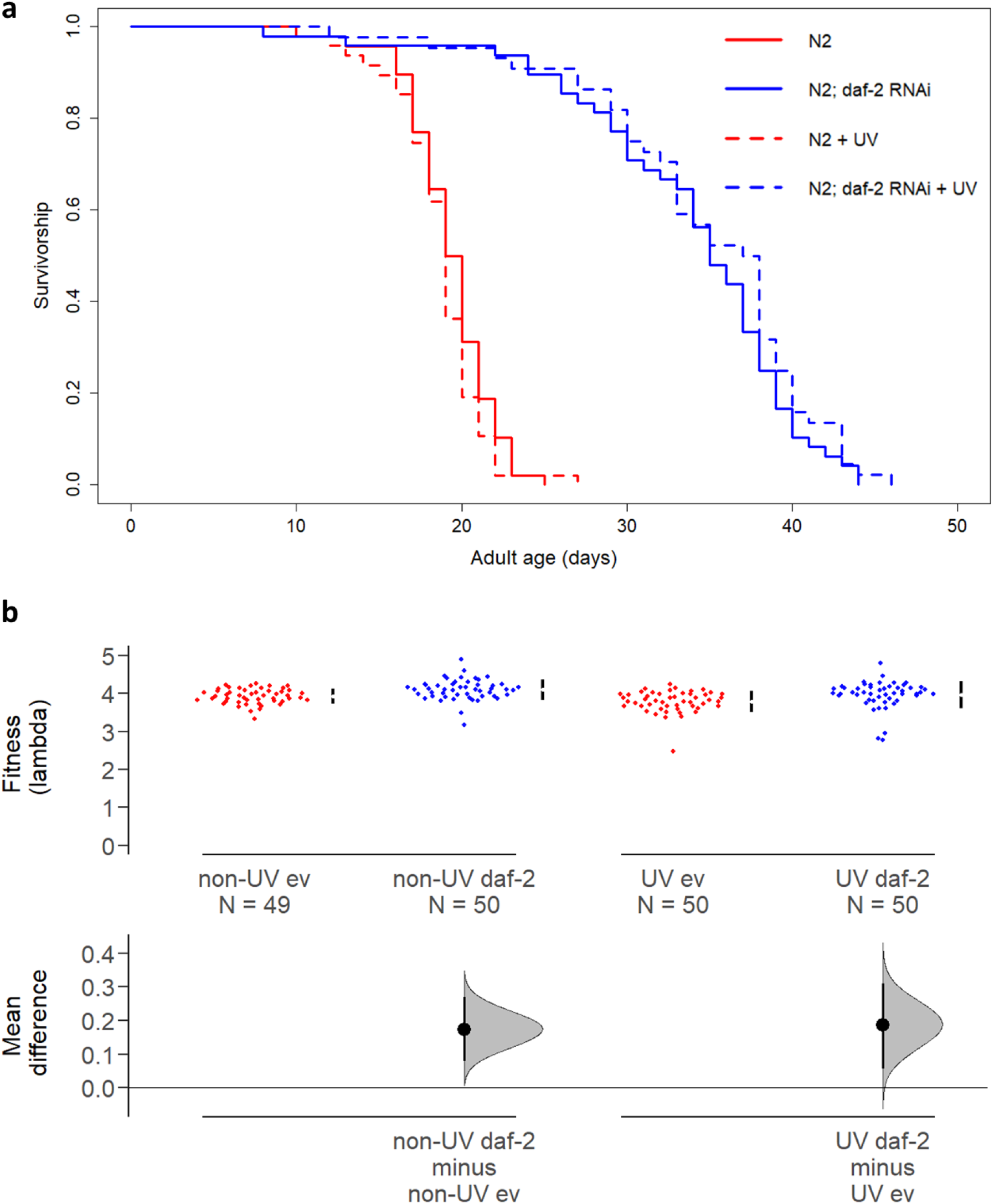
Adulthood *daf-2* RNAi in UV-irradiated and non-irradiated parents increases parental lifespan and offspring fitness. **a,** Parental survival, n=50 per treatment. **b,** Offspring fitness. UV irradiation status and adulthood RNAi treatment (either *daf-2* RNAi, ‘daf-2’, or empty vector control, ‘ev’) of parents is indicated. Offspring were maintained on empty vector and were not irradiated. Mean (effect size) and 95% confidence intervals shown were derived using non-parametric bootstrap resampling in ‘dabestr’ R package^59^.

There was no cost of *daf-2* RNAi in adulthood to parental reproduction, neither under benign conditions nor when parents were UV-irradiated (Fig. S1). Reproduction was estimated as age-specific offspring production (fecundity, Generalised Poisson, RNAi x UV x Age: z = −2.546, p=0.0109), as individual fitness (lambda, generalised linear model, GLM, RNAi: t=-0.065, df=1, p=0.948; UV: t=- 3.328, df=1, p=0.0104; RNAi x UV: t= 0.159, df=1, p=0.874) and as total lifetime reproduction (LRS) (GLM, RNAi: t=-1.887, df=1, p=0.0606; UV: t=-3.665, df=1, p<0.001; RNAi x UV: t= 0.066, df=1, p=0.948). UV irradiation did reduce parental fitness and total lifetime reproduction.

However, *daf-2* RNAi significantly increased offspring fitness both under benign conditions and when parents were UV-irradiated (lambda, GLM: RNAi: t=- 2.305, df=1, p=0.0222; UV: t=-1.625, df=1, p=0.1059; RNAi x UV: t=- 0.772, df=1, p=0.441; Fig. 1b). This absence of fitness costs for the offspring of UV-irradiated parents, despite the costs of irradiation for parental reproduction, suggests that irradiated parents may have invested in germline repair mechanisms, which improved the fitness of their offspring^18,38,66^.

The impact of parental *daf-2* RNAi and UV treatment on offspring age-specific reproduction varied with offspring age as did the number of offspring with zero fecundity (zero-inflated generalised Poisson, ZIGP, RNAi x age: z=4.864, p<0.001; UV x age: z=3.678, p<0.001; RNAi x UV: z=-0.175, p=0.861; ZI varied with age: z=- 0.227, p=0.00656 and age^2^: z=3.301, p<0.001; Fig. S2). There was no effect of parental treatment (neither *daf-2* RNAi, UV irradiation nor their interaction) on offspring total lifetime reproduction (GLM, RNAi: t=-1.141, df=1, p=0.255; UV: t=- 1.626, df=1, p=0.106; RNAi x UV: t= −0.053, df=1, p=0.958; Fig. S2).

Our findings show that under adulthood-only *daf-2* RNAi, parents increase investment into somatic maintenance resulting in increased lifespan with no survival or reproductive cost to themselves or their offspring under benign conditions validating previous work in *C. elegans*^19,20^. Importantly, we reveal that the absence of a longevity-fecundity trade-off in parents persisted under stressful conditions, when organisms have to invest into repairing UV-induced damage. Previous work in *daf-2* mutants has also shown them to be more resistant to UV irradiation^45^.

Furthermore, we show that *daf-2* knockdown in adult parents primarily influenced the timing of reproduction in their offspring. Specifically, it caused a shift to increased early life reproduction, improving offspring individual fitness, rather than an increase in total reproduction.

Timing of reproduction in both laboratory and natural *C. elegans* populations is important as *C. elegans* reproduces very fast and so the strength of natural selection (selection gradient) declines very rapidly with advancing age^67,68^. Under standard lab conditions, self-fertilising *C. elegans* hermaphrodites complete most of their reproduction within the first three days of adulthood and in nature, *C. elegans* exhibit boom-and-bust population dynamics involving periods of very rapid population growth, driven by an ephemeral food supply^68,69^. As individual fitness is heavily weighted by early reproduction and fast development (Supplementary Methods), even relatively small increases in Day 1 and Day 2 reproduction (Fig. S2) can improve individual fitness, despite having no effect on total lifetime reproduction, if later reproduction is correspondingly reduced.

Whilst an earlier study found increased total reproduction in the first generation of offspring from *C. elegans* parents treated with *daf-2* RNAi, across N2 wild-type and two other genetic backgrounds, this was also accompanied by increased offspring fitness^20^, in agreement with our results here. Under UV-induced mutagenesis, the costs of germline maintenance would be expected to become more apparent^29,38,70^. However we show that adulthood-only *daf-2* RNAi improves fitness of F1 offspring even when parents are under stress.

### Parental *daf-2* RNAi in adulthood does not reduce fitness or total reproduction under intermittent fasting

Under resource limitation via intermittent fasting (IF), there was also no cost of adulthood-only *daf-2* RNAi for individual fitness (lambda, generalised linear mixed effects model, GLMER, RNAi: t=-0.964, p=0.337; Fig. 2a). Despite a small cost of adulthood *daf-2* RNAi for fitness when fed ad libitum (AL) in this assay (GLMER, RNAi: t=2.375, p=0.019; RNAi x diet: t=-2.067, p=0.040; diet: t=- 15.595, p<0.001; Fig. 2a), there was no cost of *daf-2* RNAi for total lifetime reproduction under either IF or AL (GLMER, RNAi: t= 0.290, p= 0.772; diet: t= −5.979, p<0.001; RNAi x diet: t=-1.387, p=0.16646; Fig. 2b).

**Fig. 2.**
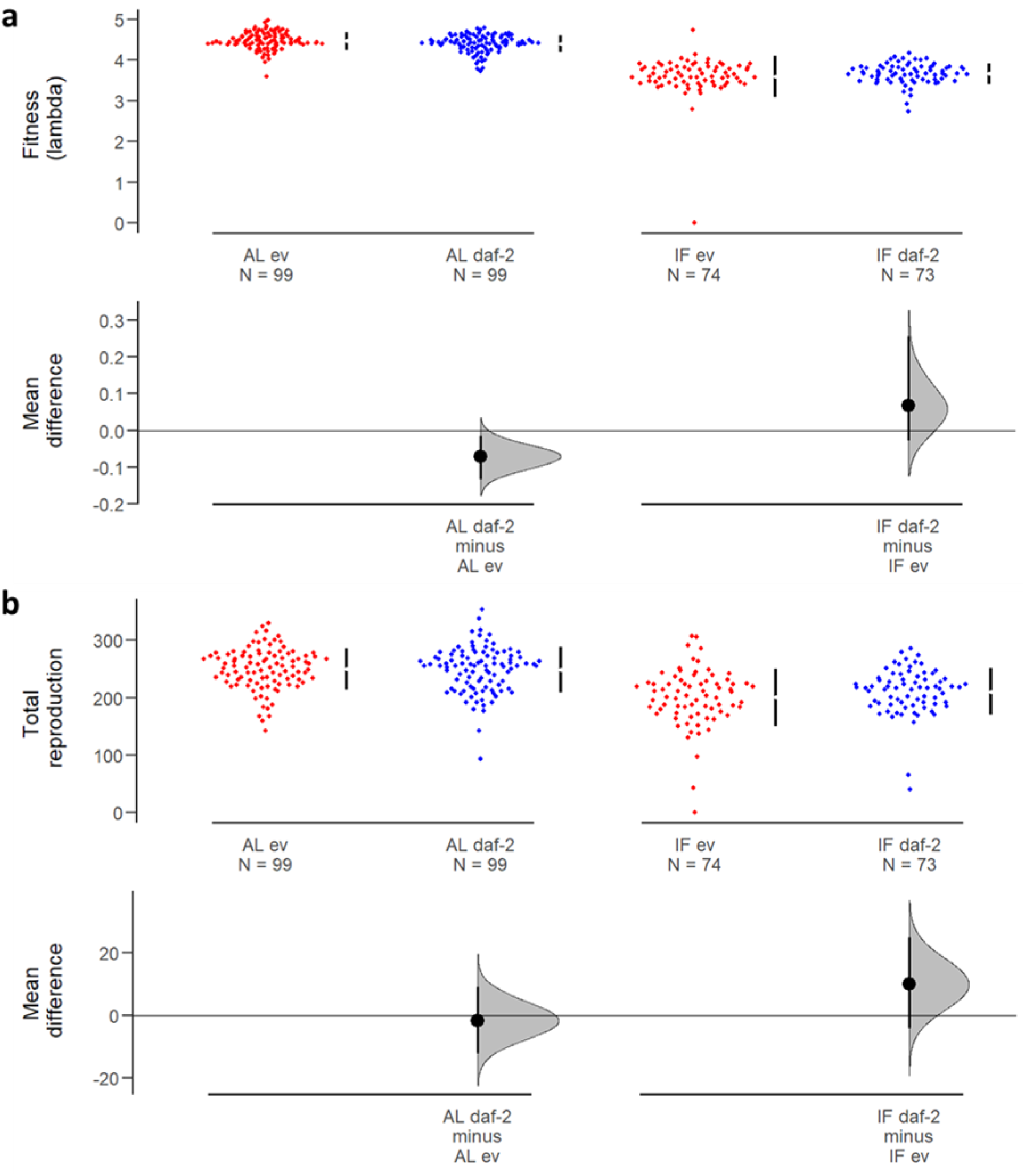
The effect of adulthood *daf-2* RNAi on parental reproduction under intermittent fasting and ad libitum feeding. **a,** Fitness (lambda). **b,** Total lifetime reproduction. Individuals were either intermittently fasted (IF) for 9h on Days 1, 3, 5 and 7 of adulthood or fed ad libitum (AL) throughout life. Adulthood RNAi treatment (either *daf-2* RNAi, ‘daf-2’, or empty vector control, ‘ev’) of parents is indicated. Adulthood *daf-2* RNAi had no effect on fitness or total reproduction under IF. Mean effect size and 95% confidence intervals shown.

We may have expected a reproductive cost of adulthood *daf-2* RNAi to become apparent under nutritional or UV irradiation stress, but instead we show no cost to fitness or total lifetime reproduction of adulthood *daf-2* RNAi under intermittent fasting (Fig. 2) or UV irradiation (Fig. S1). This is in agreement with previous work that showed *daf-2* RNAi in adult N2 *C. elegans* did not reduce lambda fitness under a variable environment of light and temperature stress^28^.

### Multigenerational *daf-2* RNAi in adulthood protects against extinction

We found that *daf-2* RNAi reduced extinction under both UV-induced and spontaneous MA across 40 generations (Cox proportional hazards regression analysis, coxph, RNAi: z=2.159, p=0.031; UV: z=11.469, p<0.001; RNAi x UV: z= −0.537, p=0.591; Fig. 3a). Extinction results did not differ significantly between two independent experimental blocks (coxph, z=-0.942, p=0.346; Fig. S3). The major causes of MA line extinction (Fig. 3b) were infertility (failure to lay eggs), or sterility (the production of eggs that did not hatch), indicative of underlying germline damage. Infertility and sterility were likely associated with observed reproductive abnormalities such as a deformed vulva, abnormal external growth close to the vulva (possible tumour) and cavities in the reproductive tract in place of embryos or oocytes (Fig. 3c). It is known that vulva-less *C. elegans* mutants are unable to lay eggs and can die from internal hatching^71,72^.

**Fig. 3.**
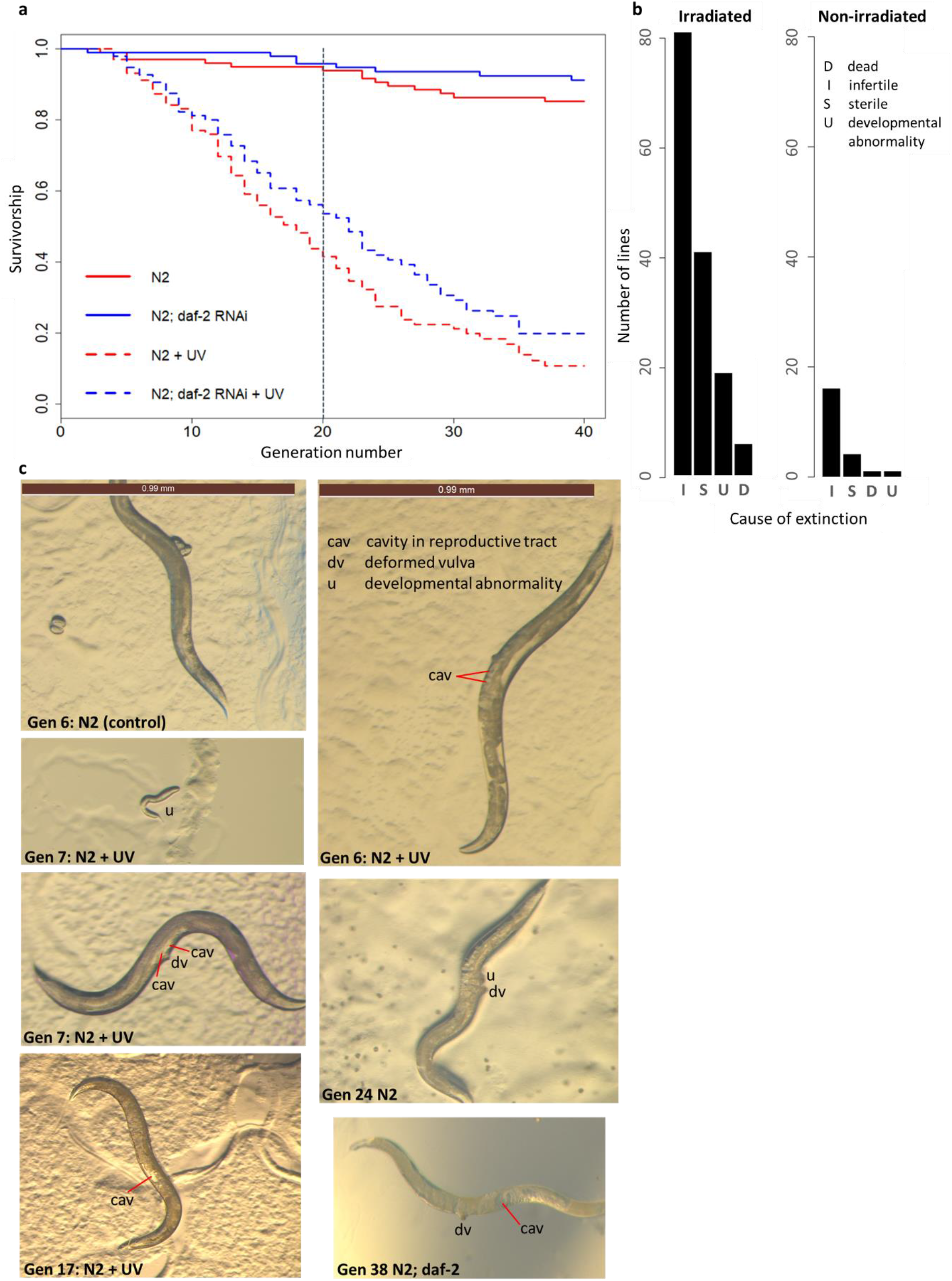
Reduced adulthood IIS, via *daf-2* RNAi, protects against N2 wild-type extinction under mutation accumulation. **a,** Transgenerational survival in N2 under spontaneous and UV-induced mutation accumulation. Vertical dotted line at generation 20 indicates timing of life history assay. Sample size of 100 lines per RNAi strain by irradiation treatment combination. **b,** Causes of extinction of N2 wild-type MA lines indicate germline damage. **c,** Representative images of germline damage. Brown scale bar of 0.99 mm for all images.

It is possible that UV irradiation may also have caused non-genetic damage (to proteins or other macromolecules) that was not inherited but may also have contributed to sterility or infertility, and that *daf-2* RNAi additionally may have protected against. Furthermore, *daf-2* RNAi may have provided enhanced stress resistance to protect against stress responses induced by UV irradiation. Despite this, previous work in *C. elegans* has shown that UV irradiation most commonly induces base substitutions, which are subject to the nucleotide excision repair pathway^73^.

The reduction in extinction observed in our MA lines with *daf-2* RNAi shows the benefits of increased investment into germline maintenance that became more pronounced across multiple generations of UV-induced and spontaneous MA.

### Multigenerational *daf-2* RNAi increased fitness of surviving MA lines

We next tested for the effects of *daf-2* RNAi on life history traits of surviving MA lines. We were interested in how IIS influences potentially detrimental effects of MA that are not sufficiently severe to cause line extinction. We assayed age-specific reproduction, egg size, male production and adult heat shock resistance in grand-offspring from the spontaneous and UV-induced MA lines on *daf-2* RNAi versus control treatments, at generation 20 of MA, following two generations of rearing under common garden conditions (no irradiation, on empty vector control) to attenuate direct effects of irradiation and RNAi. Generation 20 was a point at which there was a pronounced benefit of *daf-2* RNAi for protection against extinction in the UV-induced MA lines, but no clear difference in extinction trajectories for the spontaneous (non-irradiated) MA lines (Fig. 3a).

Adulthood-only *daf-2* RNAi across 20 generations of mutation accumulation significantly increased individual fitness in the surviving irradiated MA lines, but there was no effect of *daf-2* RNAi on the fitness of non-irradiated lines (GLM, irradiated, RNAi: t=-2.804, p=0.006; non-irradiated, RNAi: t=0.647, p=0.518; all data, RNAi x UV: t=-2.886, p=0.004; Fig. 4a), in agreement with the results on line extinction at generation 20 (Fig. 3a).

**Fig. 4.**
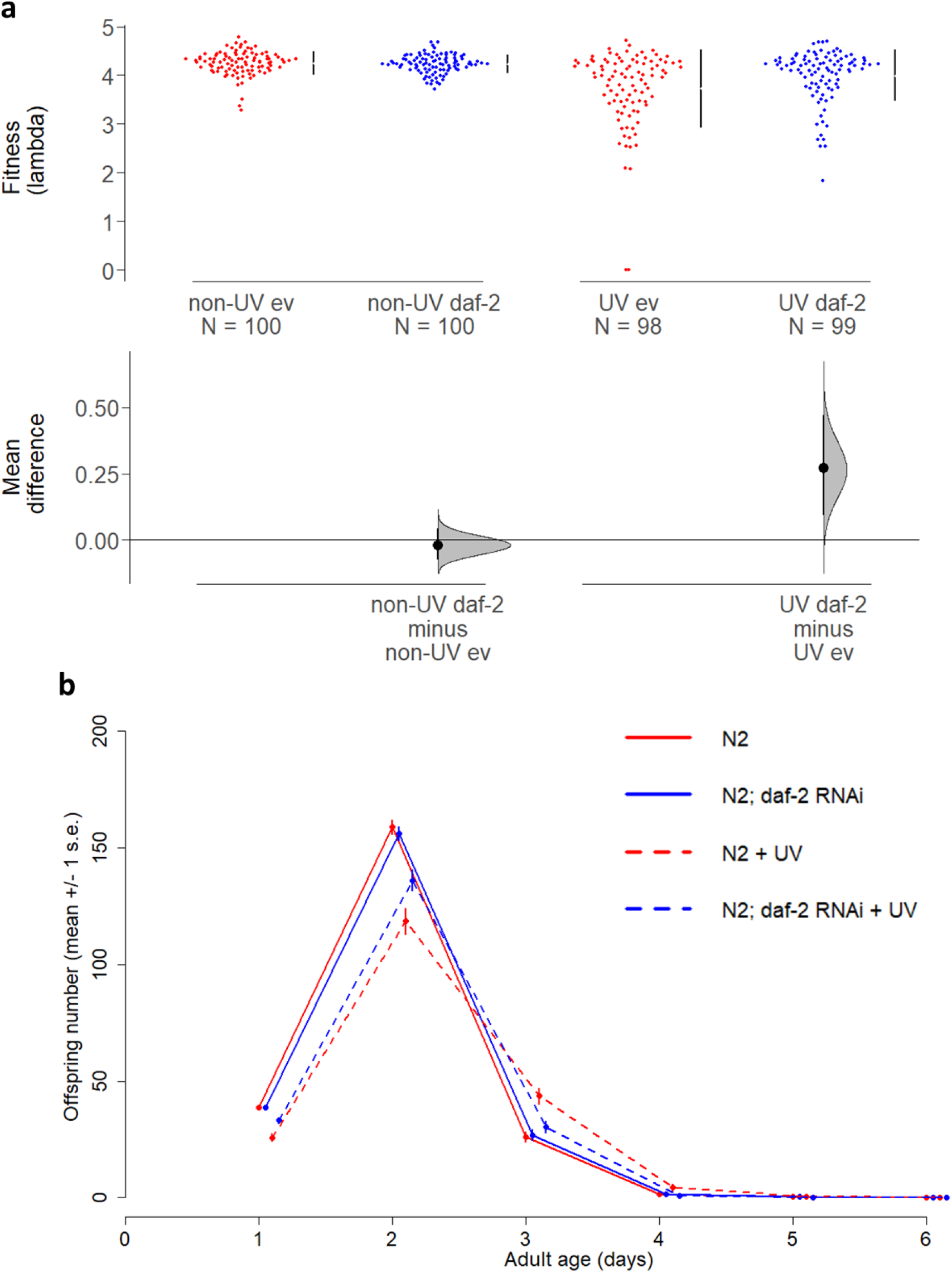
The effect of *daf-2* RNAi on fitness and age-specific reproduction after 20 generations of MA. **a,** Individual fitness (lambda) of the grand-offspring of N2 wild-type MA lines at generation 20. Fitness was assayed in a standard common garden environment, on the empty vector control and no irradiation; following two generations of rearing under standard conditions, from MA generation 20. Mean and 95% confidence intervals shown. **b,** Age-specific reproduction in the grand-offspring of N2 MA lines at generation 20.

The increased fitness of irradiated *daf-2* RNAi MA lines was driven by their improved early life fecundity (Day 1 and Day 2 offspring production), relative to irradiated controls (ZIGP, RNAi x age: z=-1.981, p=0.0475; RNAi x age^2^: z=3.334, p<0.001; Fig. 4b), an effect absent in the non-irradiated lines (ZIGP, non-UV, RNAi: z=0.05, p=0.963; all data: RNAi x UV x age: z=-2.50, p=0.0125; RNAi x UV x age^2^: z=3.58, p<0.001; ZI intercept: z=-18.27, p<0.001). There was no effect on total reproduction (GLM, UV: t=-0.811, p=0.419; non-UV: t=0.875, p=0.383; UV x RNAi: t=-1.138, p=0.256; Fig. S4).

We suggest that the fitness benefits of *daf*-2 RNAi for irradiated MA lines were most likely due to genetic differences between the treatments, as the benefits persisted after two generations of common garden rearing and therefore were neither due to the direct exposure of offspring to RNAi nor acute effects of UV (neither as adults nor as eggs). This conclusion is reinforced by the finding that parental *daf*-2 RNAi effects on offspring fitness do not persist beyond F1 (see below, Fig. 6). Variation in fitness was considerably greater across irradiated than across non-irradiated MA lines (Fig. 4a), suggesting that irradiation-induced *de novo* mutations generated greater genetic and phenotypic variation compared to spontaneous MA. The spontaneous mutation rate in N2 wild type *C. elegans* is estimated as one *de novo* mutation per individual per generation, under standard conditions^73,74^. The close association between extinction trajectories during the first 20 generations of MA (Fig. 3a) and the fitness of the surviving MA lines (Fig. 4a) suggests that a threshold of accumulating deleterious mutations needs to be crossed before the effect size is sufficiently severe to result in extinction.

The most parsimonious explanation for the reduced extinction and increased fitness of *daf-2* RNAi irradiated lines would therefore be due to genetic differences in the rate of MA. To directly distinguish between genetic and non-genetic effects associated with MA line extinction, future work could use whole genome sequencing (WGS) to quantify mutation rates between the MA treatments.

To determine if increased offspring fitness in the surviving *daf-2* RNAi-treated MA lines, was associated with greater parental resource allocation into their eggs, we measured egg size, as a proxy for parental investment (n=75 eggs measured, one per individual taken from the Generation 20 fitness assay). Previous work has shown that reduced IIS, either via dietary restriction or via *daf-2* RNAi, increases mean egg size^20,57^. Egg size has been correlated with offspring fitness^20^ and larval body length^75^ in *C. elegans*.

We found that grandparental *daf-2* RNAi resulted in grand-offspring (F2) that laid smaller eggs if their grand-parents from the MA lines had been irradiated, but there was no significant effect on the size of eggs laid by F2 offspring descended from non-irradiated grandparents treated with *daf-2* RNAi following 20 generations of MA (GLM, UV lines, RNAi: t=2.362, df=1, p= 0.0195; non-UV lines, RNAi: t=-1.246, df=1, p=0.215; all data, RNAi x UV: t=2.642, df=1, p=0.00869; RNAi x UV x Block: t=- 0.774, df=1, p=0.440; Fig. 5). This is contrary to the increase in F1 egg size under reduced parental IIS found in previous work under benign conditions^20,57^.

**Fig. 5.**
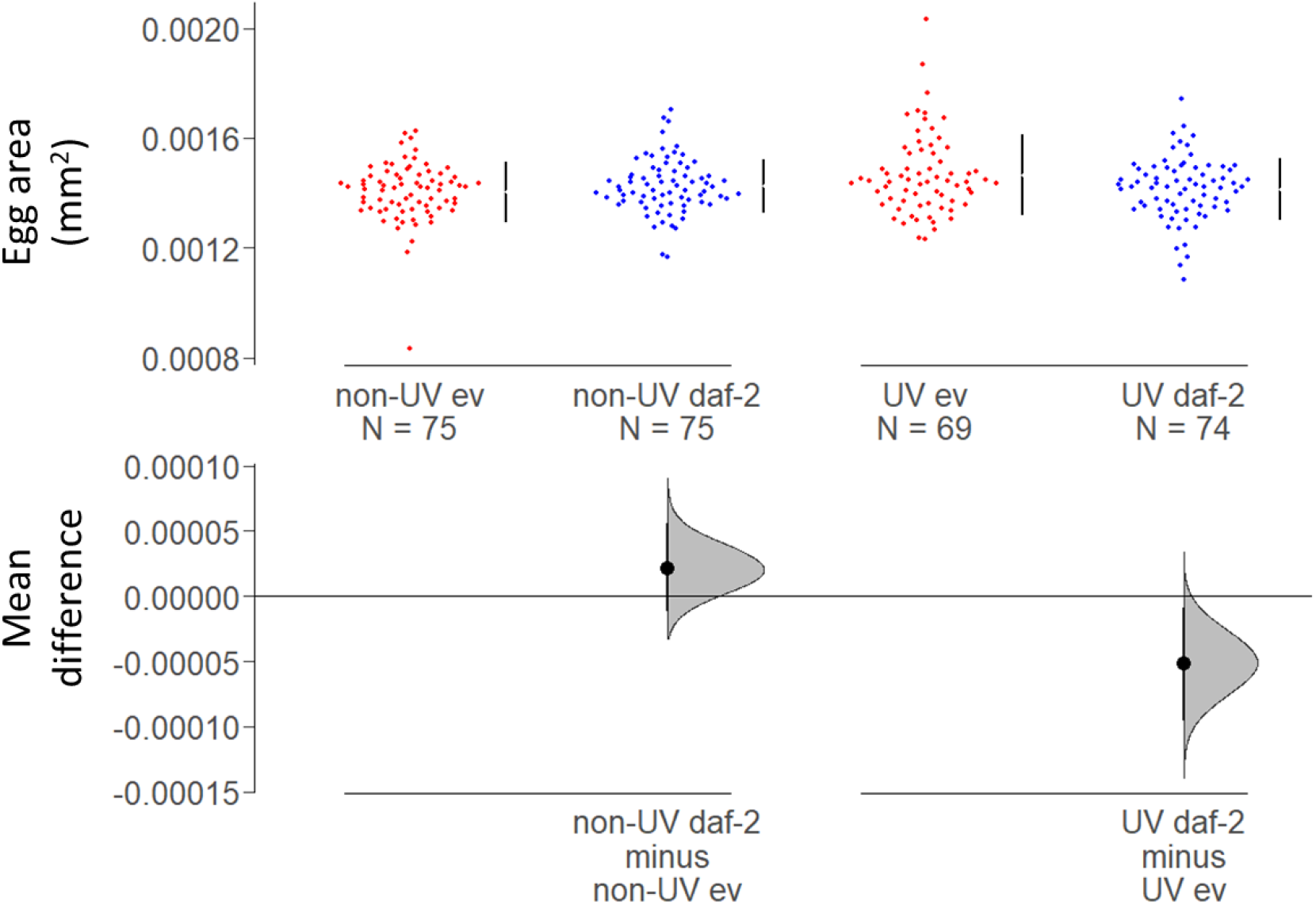
The effect of *daf-2* RNAi on egg size after 20 generations of spontaneous or UV-induced MA. Egg size (area, mm^2^) was reduced in the grand-offspring of UV-induced MA lines on *daf-2* RNAi (‘daf-2’) relative to those from UV-irradiated MA lines on empty vector control (‘ev’). However, this difference in egg size was absent from the spontaneous (‘non-UV’) MA lines, following 20 generations of MA. Grand-offspring were all non-irradiated and kept on ev. Mean and 95% confidence intervals are shown.

Our results indicate that even though F2 offspring from irradiated *daf-2* RNAi MA lines laid smaller eggs, this did not seem to impact negatively on the quality (fitness) of offspring they produced. It is possible that a smaller egg size could be a phenotypically plastic response to UV radiation, perhaps to improve stress resistance to irradiation. Alternatively, it could be the result of a trade-off between improved investment into germline genetic quality and egg size. At present, it is unclear why grandparental *daf*-2 RNAi resulted in smaller eggs in F2.

Exposure to certain environmental stresses (e.g. high temperature^76^, starvation^77^, increased mutation rate^78^), increases male production and outcrossing in *C. elegans*. However, we found no increase in the proportion of males produced by the grand-offspring in irradiated MA lines. Only one male developed from the 293 eggs assayed.

Reduced IIS increases the activity of heat shock factor 1 which mediates lifespan extension^79,80^. Heat shock responses are conserved across diverse taxa and positively associated with lifespan (as reviewed by^81,82^). To test the stress resistance of post-reproductive (Day 7) adults derived from the Generation 20 MA lines, we assayed survival and locomotion under acute heat shock for 1 hour and 45 minutes at 37°C (following^83^). We found no effect of MA line treatment (neither UV irradiation nor *daf-2* RNAi) on the survival of untreated Day 7 F2 offspring following heat shock. Only 4% of the 371 individuals were dead by 24 hours after heat shock. This high post-heat-shock survival of Day 7 adults is in line with previous work, which found survival after heat shock to more than double with age, from the first to the fourth day of adulthood in *C. elegans^83^*. However F2 offspring from non-irradiated *daf-2* RNAi MA lines recovered normal locomotion faster following heat shock than F2 from non-irradiated MA control lines, but this benefit was not seen in descendants from irradiated MA lines (Binomial GLM, 3h post-heat shock, RNAi: z=-2.274, df=1, p=0.0229; UV: z=-2.078, df=1, p=0.0377; RNAi x UV: z=2.304, df=1, p= 0.0212; 24h post-heat shock, UV: z=-0.206, df=1, p=0.837; RNAi: z=-0.045, df=1, p=0.964; RNAi x UV: z=1.347, df=1, p=0.178; Fig. S5).

### Parental *daf-2* RNAi does not affect offspring fitness transgenerationally

Transgenerational epigenetic inheritance of RNAi can occur in *C. elegans* via transmission of small interfering RNAs^43,84^. This transgenerational inheritance of RNAi can last for several generations^84–86^ and requires the germline argonaut protein, HRDE-1, that is absent in *hrde-1* (heritable RNAi defective-1) mutants^87^. RNAi is still functional in somatic cells, as the *hrde-1* mutation only affects the inheritance of RNAi via the germline. To determine whether there was direct transgenerational transfer of *daf-2* RNAi via the germline and thus for how many generations the effects of parental *daf-2* RNAi could persist, we assayed the fitness effects on three generations of untreated offspring (F1, F2 and F3) from *daf-2* RNAi treated versus control parents. Two genetic backgrounds were used: *C. elegans* N2 wild-types, and *hrde-1* mutants that did not transfer RNAi transgenerationally.

Effects of parental *daf-2* RNAi on the fitness of untreated descendants were absent after the first generation of offspring in the N2 wild-types and absent across all generations of offspring in the *hrde-1* mutants (GLM, F1, RNAi x genotype: t=- 2.160, df=1, p=0.0329; F2, RNAi x genotype: t=0.372, df=1, p=0.711; RNAi: t=1.090, df=1, p=0.278; genotype: t=7.044, df=1, p<0.001; F3, RNAi x genotype: t=- 0.973, df=1, p=0.332; RNAi: t=0.359, df=1, p=0.721; genotype: t=5.302, df=1, p<0.001; all data: RNAi x generation: F=4.450, df=2, p=0.0124; Fig. 6). Furthermore, there was no effect of parental *daf-2* RNAi on the age-specific reproduction of offspring beyond F1, for either N2 or *hrde-1* backgrounds (Fig. S6; Table S2). There was also no parental *daf-2* RNAi effect on total reproduction for F2 and F3 offspring generations (Fig. S7; Table S2).

**Fig. 6.**
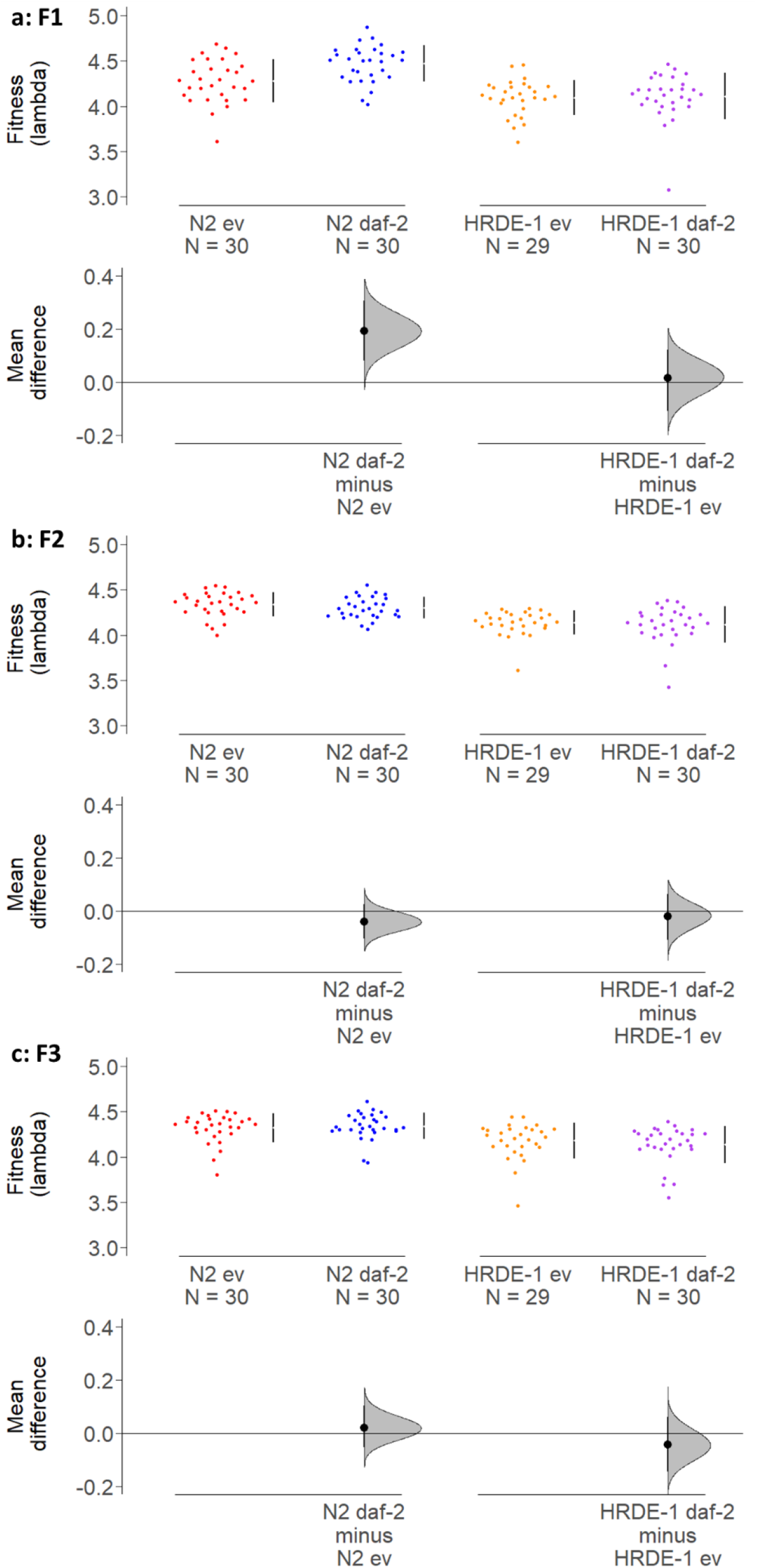
The effects of *daf-2* RNAi on offspring fitness do not persist beyond the first offspring generation. Fitness of the: **a,** first (F1); **b,** second (F2) and **c,** third (F3) generation of offspring from parents treated with *daf-2* RNAi (‘daf-2’) or an empty vector control (‘ev’) as indicated, in N2 wild-type and RNAi inheritance deficient *hrde-1* mutant backgrounds. All offspring generations were untreated (kept on ev).

The absence of fitness benefits in the second and third generations of offspring from *daf*-2 RNAi parents, implies that *daf-2* RNAi is not transgenerationally inherited beyond the first generation of offspring in the N2 background. This strongly suggests that the life-history differences between the *daf-2* RNAi and control irradiated MA lines at generation 20 in the common garden experiment were due to genetic differences and not the direct effects of inherited RNAi. Interestingly, the heritable RNAi deficiency 1 (*hrde-1*) gene was required for the fitness benefits of parental *daf-2* RNAi in F1 offspring, as these offspring fitness benefits were absent in the *hrde-1* mutant background. The *hrde-1* mutants had overall lower fitness and total reproduction than N2 wild-type.

### *hrde-1* is required for *daf-2* RNAi to confer germline protection under MA

*hrde-1* encodes a germline Argonaut protein that plays a key role in nuclear RNAi, RNAi inheritance and promoting germline immortality^88^. To determine if functional *hrde-1* was necessary for the protective effects of *daf-2* RNAi under spontaneous and UV-induced MA, we ran 400 MA lines in parallel to the N2 MA experiment, using the *C. elegans hrde-1* mutant background and reduced IIS, via *daf-2* RNAi, in half of the MA lines. The heritable RNAi deficiency (*hrde-1* mutant) resulted in the rapid extinction of irradiated MA lines and the loss of the protective effects of *daf-2* RNAi under UV-induced MA, across 25 generations (RNAi: z=-4.016, p<0.001; UV: z=12.370, p<0.001; RNAi x UV: z=-1.758, p=0.079; Fig. 7). In fact, irradiated *daf-2* RNAi lines went extinct faster than controls, in the *hrde-1* mutant background. This represented a significant difference between *hrde-1* mutant and N2 wild type backgrounds in the effect of *daf-2* RNAi (although not UV) on MA line extinction (RNAi x genotype: z=4.480, p<0.001; genotype: z=-8.083, p<0.001; RNAi: z=-4.351, p<0.001; UV x genotype: z=-0.487, p=0.626; UV: z=17.148, p<0.001).

**Fig. 7.**
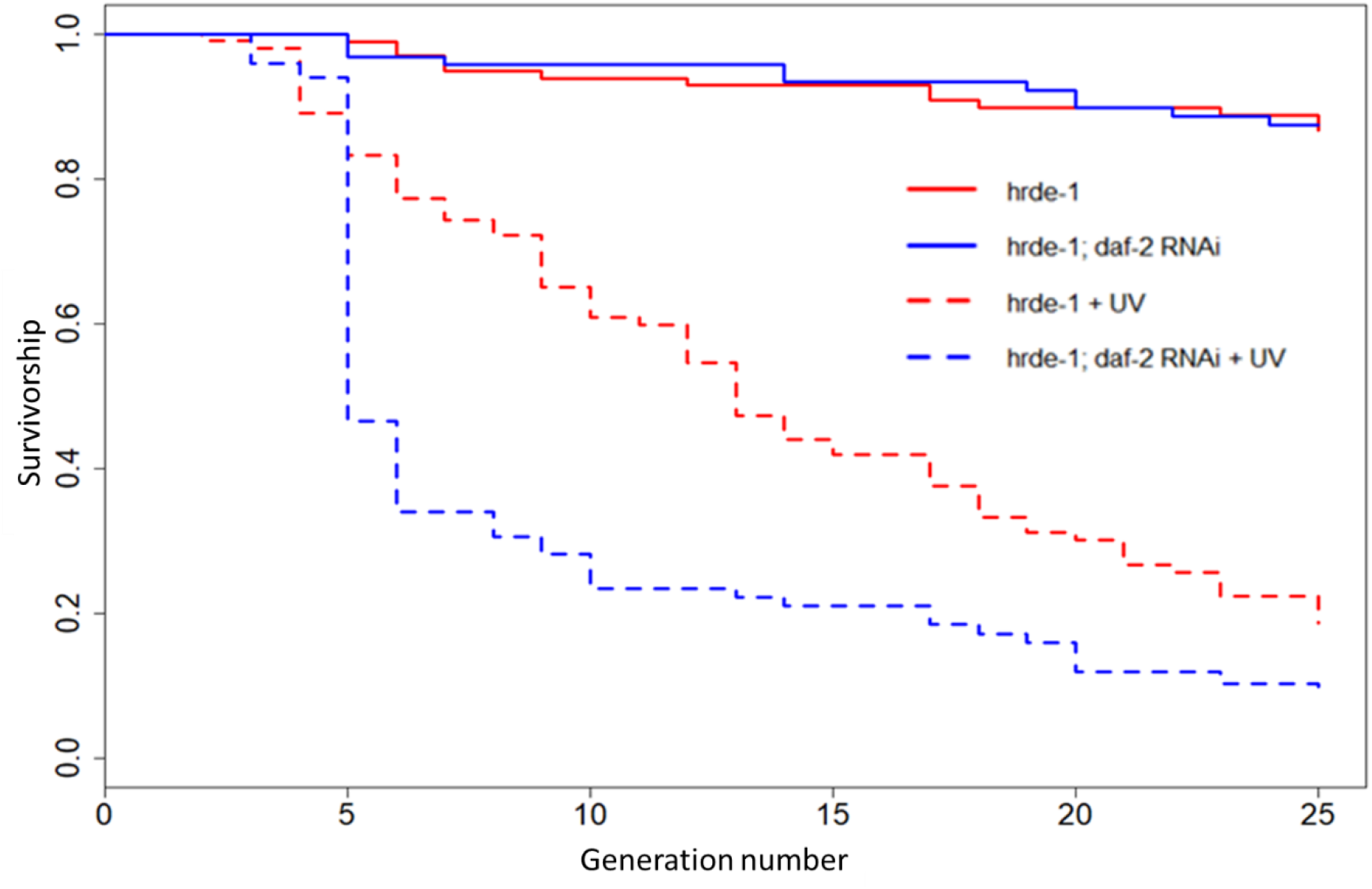
The effect of *daf-2* RNAi on *hrde-1* mutant multigenerational survival under mutation accumulation. Sample size of 100 MA lines for each RNAi by UV irradiation treatment combination.

The major cause of extinction in irradiated *hrde-1* mutant MA lines was developmental arrest, followed by infertility (Fig. S8). This strongly suggests that functional RNAi, mediated by the HRDE-1 germline argonaut protein, is required for *daf-2* RNAi protection against extinction, and for normal development and reproduction under UV-induced MA.

We show the important role of *hrde-1* in the *daf-2* RNAi-mediated protection of irradiated offspring under mutation. Our results suggest that the interaction between UV-induced germline damage and a deficiency in transgenerational inheritance of RNAi reverses the protective benefits of *daf-2* RNAi. This supports previous work suggesting that *hrde-1* mutants suffer from progressive sterility under high temperatures, implying their increased sensitivity to environmental stresses driven in part by defects in gametogenesis^87^.

### Multigenerational *daf-2* RNAi in adulthood downregulates *daf-2* expression in MA lines at generation 20

We confirmed, using qRT-PCR analysis, that feeding nematodes bacteria that expresses double-stranded RNA for *daf-2* significantly downregulated *daf-2* expression in the MA lines at generation 20 (Table S3). The extent of *daf-2* downregulation at generation 20 varied between the MA treatments (Table S3; Fig. S9). For N2 wild types, *daf-2* expression was downregulated by 23% in UV-induced MA lines and by 12% in spontaneous MA lines (based on mean gene expression fold change, 2^-ΔΔCT^). For *hrde-1* mutants under spontaneous MA, *daf-2* expression was more subtly downregulated (by 6% fold change; Table S3; Fig. S9). We were unable to analyse data from UV-induced MA *hrde-1* mutant lines as they did not recover well from freezing (only n=1 MA line for the *daf-2* RNAi treatment; Fig. S9).

The degree of downregulation of *daf-2* expression between MA treatments, interestingly matched the different extent of life history divergence between *daf-2* RNAi and empty vector treatments for the respective MA treatments at generation 20, in both N2 and *hrde-1* mutant extinction trajectories (Fig. 3a; Fig. 7) and in N2 fitness (Fig. 4a). The relatively modest degree of *daf-2* downregulation in the spontaneous MA lines for N2, and particularly for *hrde-1* mutants, in comparison to the 50% knockdown of *daf-2* expression from adulthood *daf-2* RNAi found in previous work on Day 2 N2 adults^28^, could be due to an interaction between RNAi knockdown and genetic background, or a possible ‘acclimatisation’ to the multigenerational application of *daf-2* RNAi in adulthood, but the exact reasons would require further investigation.

Technical replicates were highly repeatable for all samples (CV < 2.4% and CV < 1.5% for 98% of samples). Biological replicates showed more variation in relative gene expression within each MA line (mean CV= 7%; max. CV =8%), but within the expected range for *C. elegans* based on previous qRT-PCR analyses for different genes^89,90^.

## Conclusion

We found that reduced insulin signalling in adulthood, via *daf-2* RNAi, protects against extinction under both UV-induced and spontaneous mutation accumulation in *C. elegans*. Furthermore, the fitness of the surviving UV-irradiated MA lines was higher under *daf-2* RNAi. Most extinctions occurred because of infertility, egg hatching failure and developmental abnormalities suggesting that germline mutation accumulation directly contributed to the observed differences between the RNAi treatments. Future assessment of mutation rates in the MA lines could be useful to distinguish between genetic and non-genetic effects on extinction. Germline protection under *daf-2* RNAi requires nuclear argonaut *hrde-1* because fitness of *hrde-1; daf-2* RNAi worms was reduced both in one-generation and in multi-generation experiments. This is in line with previous work suggesting that *hrde-1* is required for transgenerational inheritance and germline immortality^87,88^.

We set out to test whether adulthood-only *daf-2* RNAi, known to extend lifespan without an obvious cost to parental fecundity^19,20^, trades-off with detrimental fitness effects when applied across multiple generations. However, we found that multigenerational downregulation of *daf-2* via adulthood-only RNAi has positive effects on the germline and protects mutation accumulation lines from extinction. This was particularly so when germline mutation rate was increased by low-level UV radiation. Our results therefore suggest that while wild-type levels of *daf*-2 expression promote development, growth and early-life reproduction, they are costly for late-life fitness and this cost accumulates across generations leading to lineage extinction.

Antagonistic pleiotropy theory of ageing (AP) maintains that genes that increase fitness in early life at the expense of fitness late in life can be overall beneficial for fitness and go to fixation^6^. In line with AP, downregulation of *daf-2* expression during development reduces fitness^19,41^, but what is the physiological basis of the putative trade-off? Our findings that adulthood-only downregulation of *daf-2* expression protects both the germline and the soma under benign conditions and under UV-induced stress, argue against the idea that resource allocation underlies this trade-off. However, our results are in line with the hypothesis that selection optimises physiological processes for development and early-life reproduction but fails to optimise the regulation of gene expression associated with these same physiological processes later in life^2,10,22–24^. This is likely because selection on insulin signalling in adulthood is too weak in *C. elegans^91^*. Future studies should focus on investigating the contribution of weakening selection on gene expression with age to the evolution of ageing across taxa, on testing the effects of downregulated IIS on survival and fitness under wide range of ecologically relevant stresses, and on developing a comprehensive theoretical framework that links genetic and physiological theories of ageing.

