## Supplementary Material for "Multigenerational downregulation of insulin/IGF-1 signalling in adulthood improves lineage survival, reproduction, and fitness in *C. elegans* supporting the developmental theory of ageing"

**Supplementary Figures and Tables**

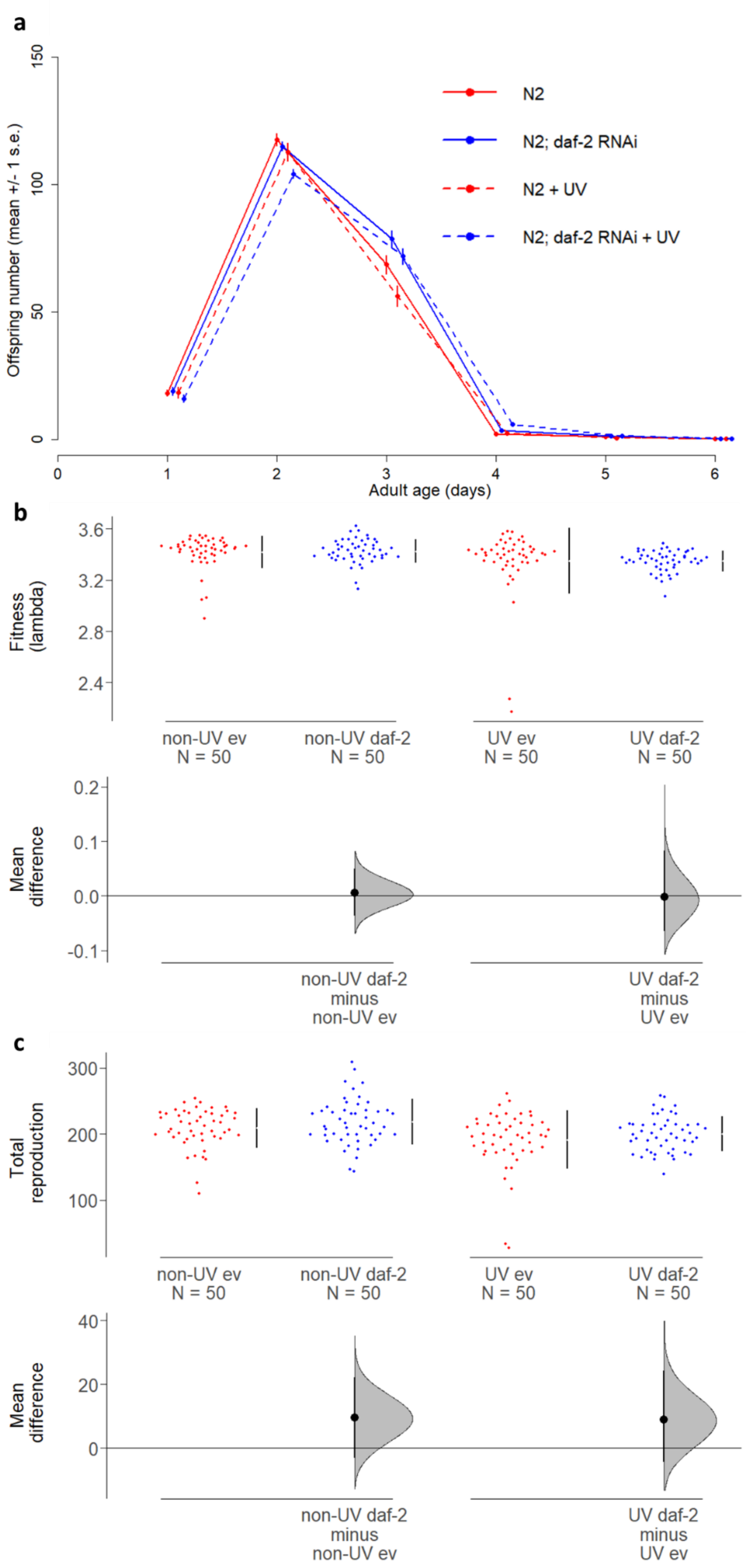


**Supplementary Figure 1: No effect of *daf-2* RNAi in adulthood on parental reproduction. a,** Age-specific reproduction. **b,** Fitness (lambda). **c,** Total reproduction. Mean and 95% confidence intervals shown.

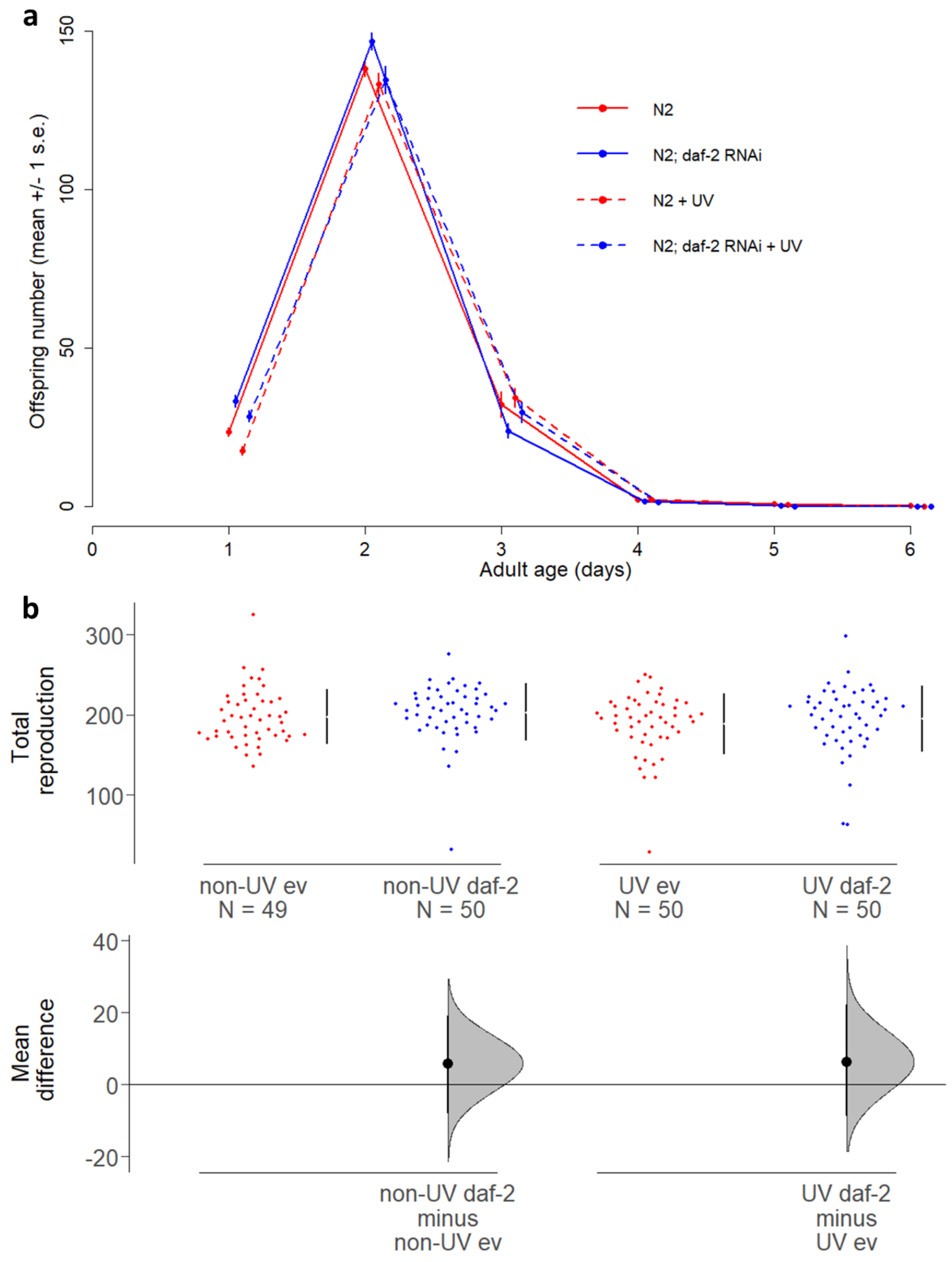


**Supplementary Figure 2: The effect of *daf-2* RNAi in UV-irradiated adult parents, on the reproduction of their offspring. a,** Age-specific reproduction. **b,** Total reproduction. Parents were UV-irradiated or not and on *daf-2* RNAi ('daf-2') or an empty vector control ('ev'). Parental *daf-2* RNAi affected offspring age-specific reproduction (when parents were irradiated), but not offspring total reproduction.

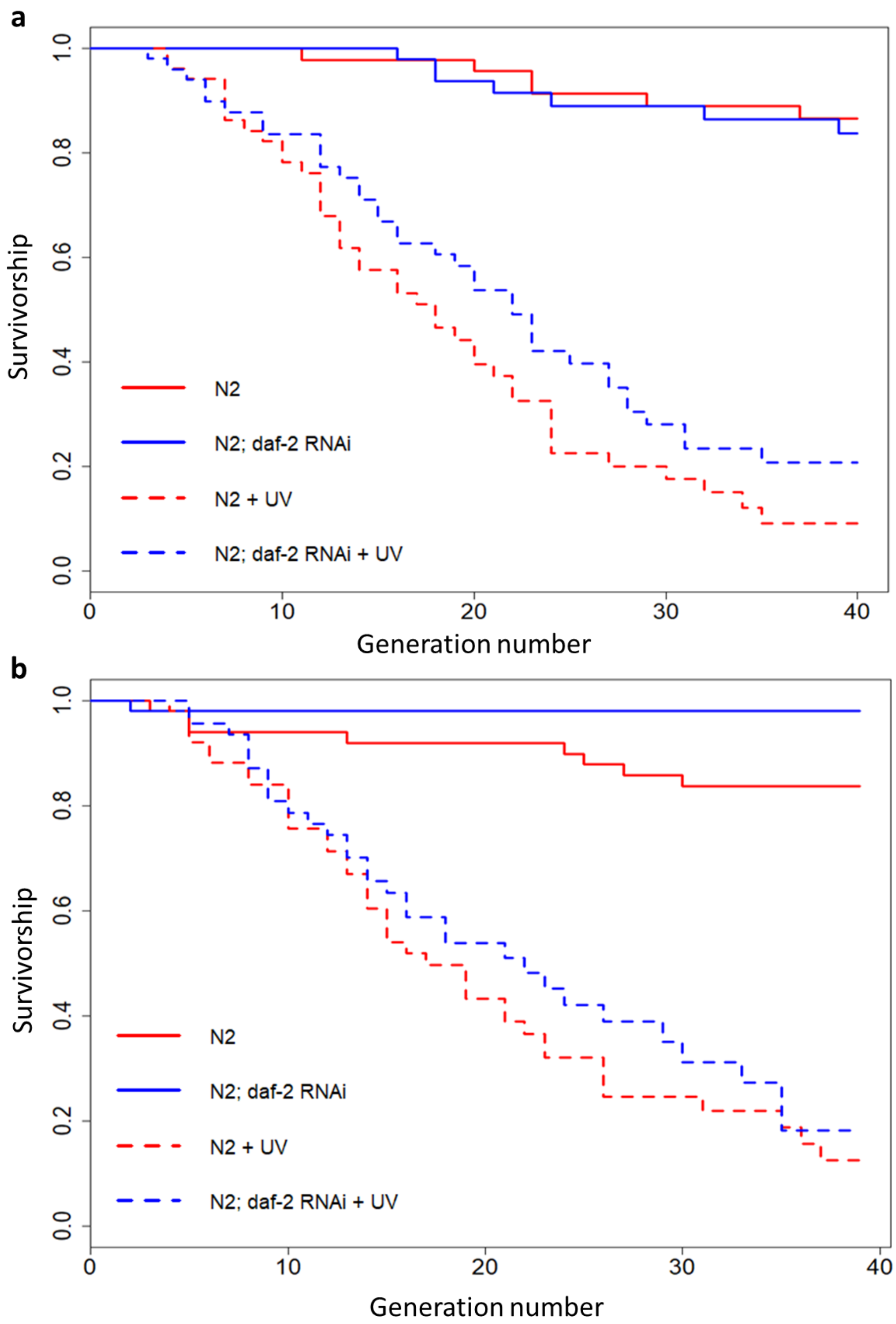


**Supplementary Figure 3: Transgenerational survival under mutation accumulation in wild-type (N2) background, by block. a,** Block 1. **b,** Block 2. Sample size of 50 MA lines per treatment per block.

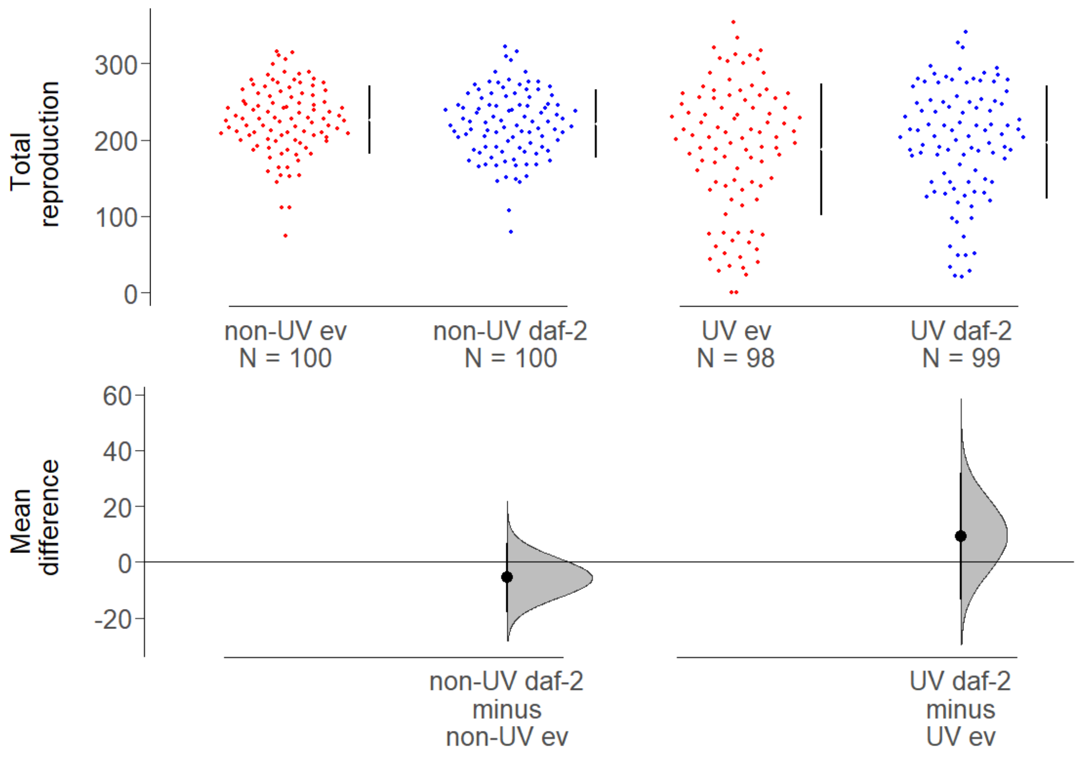


**Supplementary Figure 4: No difference in total reproduction of surviving MA lines at generation 20 in wild-type (N2) background.** Total lifetime reproduction of MA lines was calculated from the sum of daily reproduction in a standard common garden environment, on the empty vector control and no irradiation; following two generations of rearing under standard conditions, from MA generation 20. Treatments of MA lines indicated.

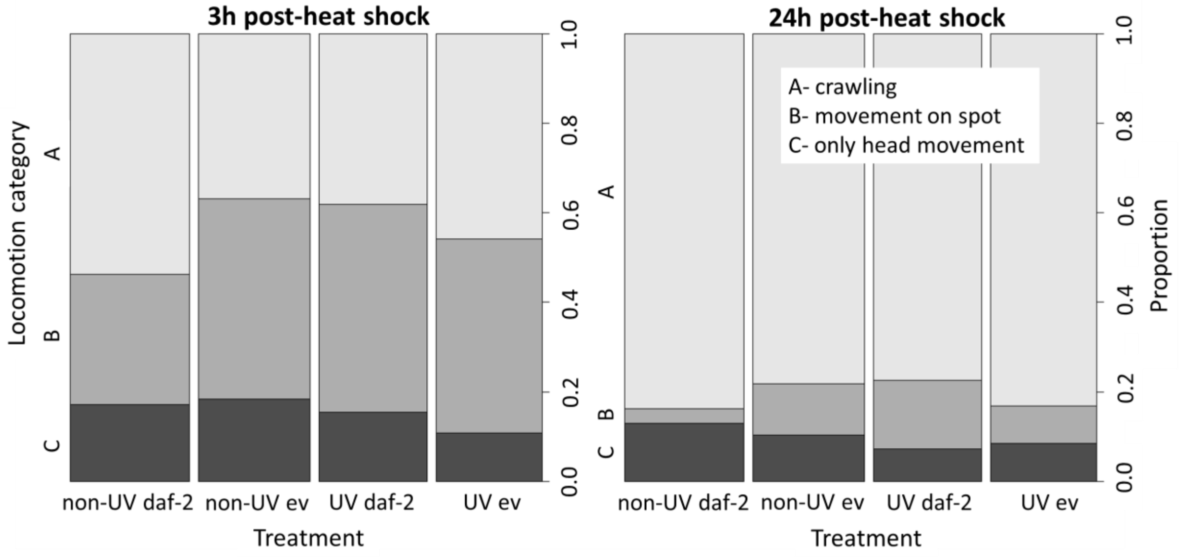


**Supplementary Figure 5: Effect of heat shock on locomotion of Day 7 adults in surviving MA lines at generation 20 in wild-type (N2) background.** Locomotion (a measure of heat shock resistance) was recorded in the same individuals at 3 hours and 24 hours after the end of heat shock (37°C for 1 hour 45 minutes). Same individuals as used in the reproduction assay of MA lines from generation 20. Treatments of MA lines indicated (n= 93, 87, 84, 83 individuals per treatment, L-R respectively). The few individuals that died from heat shock were excluded from locomotion analysis.

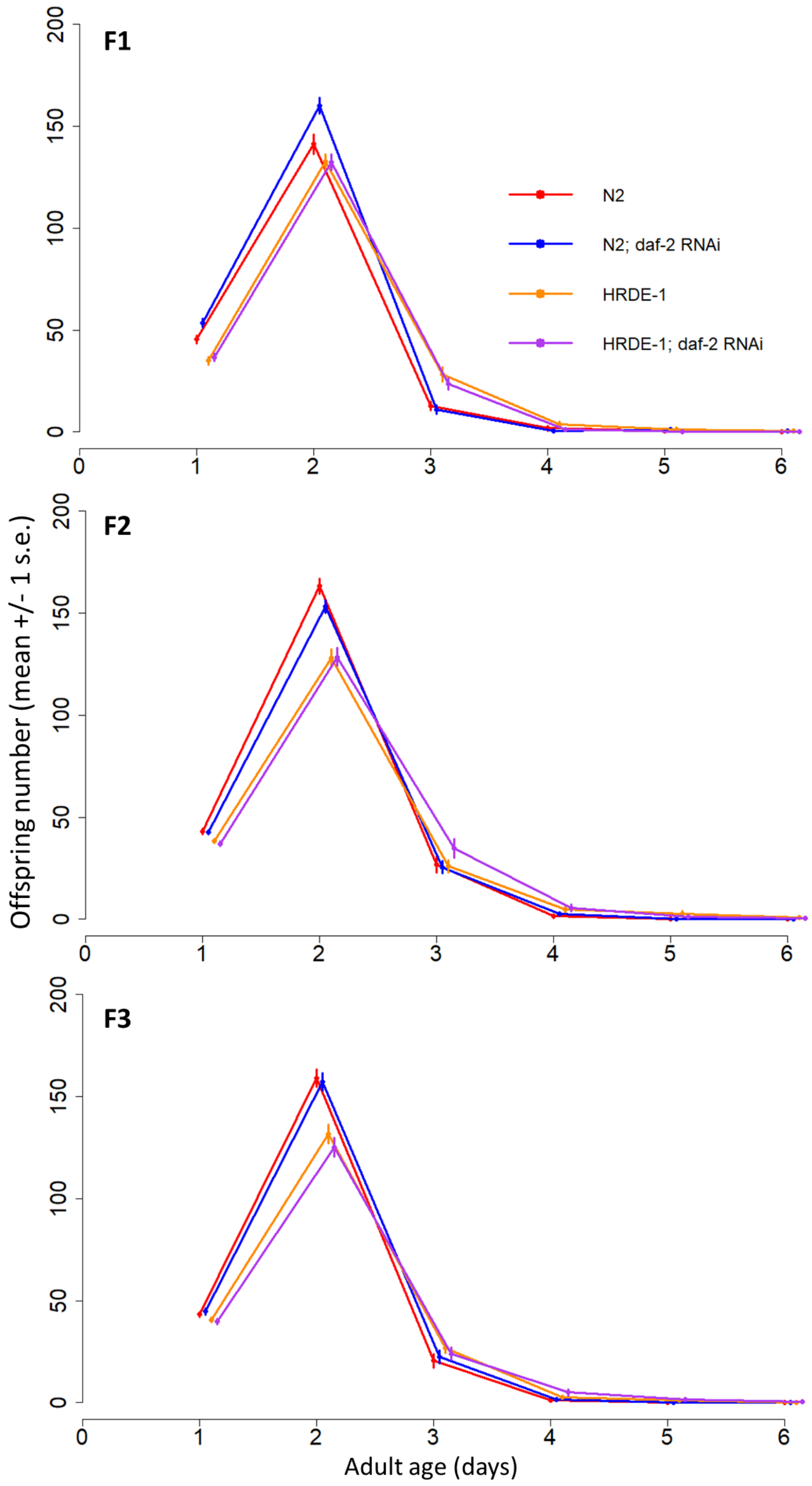


**Supplementary Figure 6: The effects of *daf-2* RNAi on offspring age-specific reproduction do not persist beyond the first offspring generation.** Reproduction counts for the first (F1), second (F2) and third (F3) generation of offspring from parents treated with *daf-2* RNAi ('daf') or an empty vector control ('ev'), in N2 wild-type and RNAi inheritance deficient *hrde-1* mutant backgrounds. All offspring generations were kept on ev. Sample sizes: N2 (n=30), N2; *daf-2* RNAi (n=30), *hrde-1* (n=29) and *hrde-1*; *daf-2* RNAi (n=30).

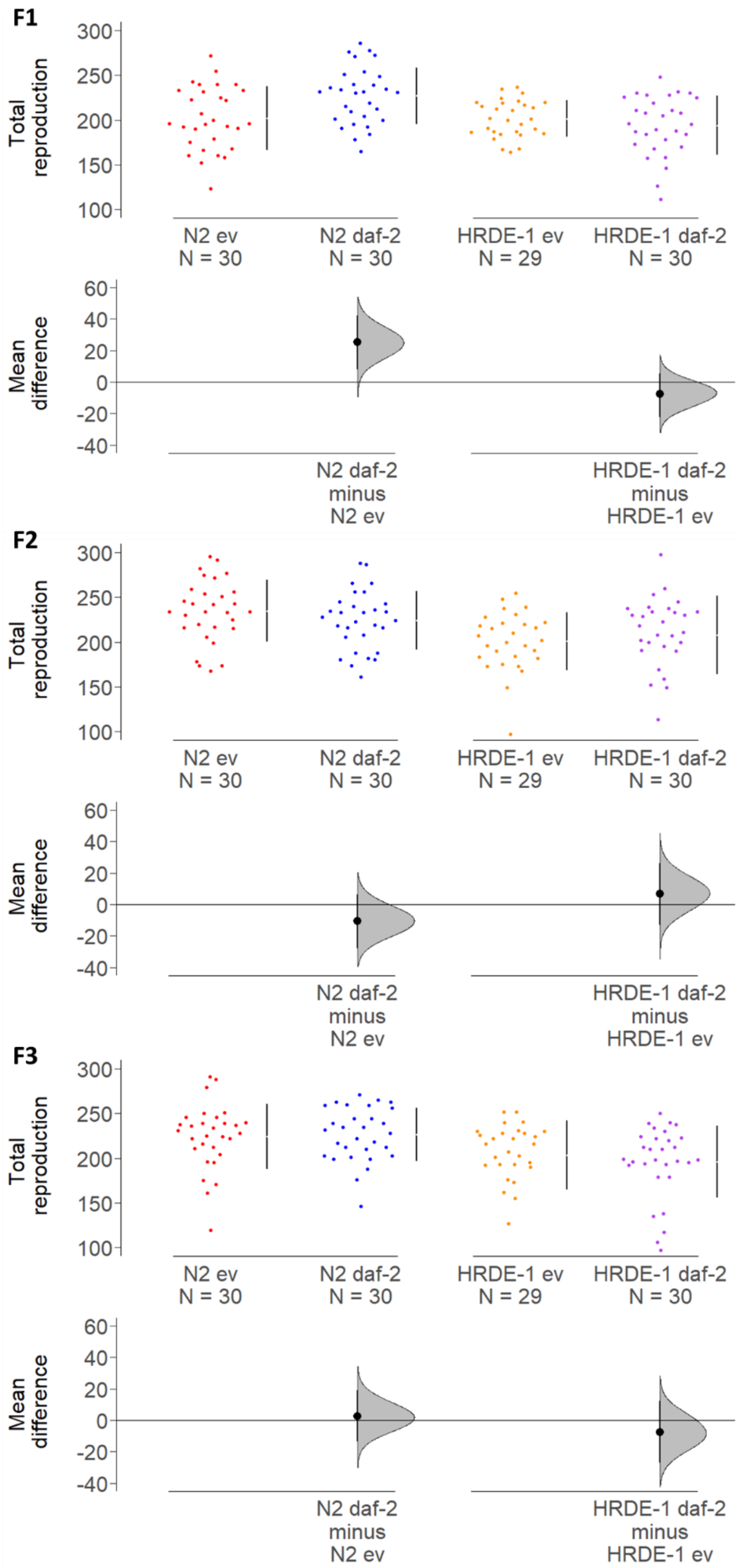


**Supplementary Figure 7: The effects of *daf-2* RNAi on offspring total lifetime reproduction do not persist beyond the first offspring generation.** Total lifetime reproduction counts are the sum of age-specific reproduction counts for each individual, in the first (F1), second (F2) and third (F3) generation of offspring from parents treated with *daf-2* RNAi ('daf-2') or an empty vector control ('ev'), in N2 wild-type and RNAi inheritance deficient *hrde-1* mutant backgrounds. All offspring generations were untreated (kept on ev).

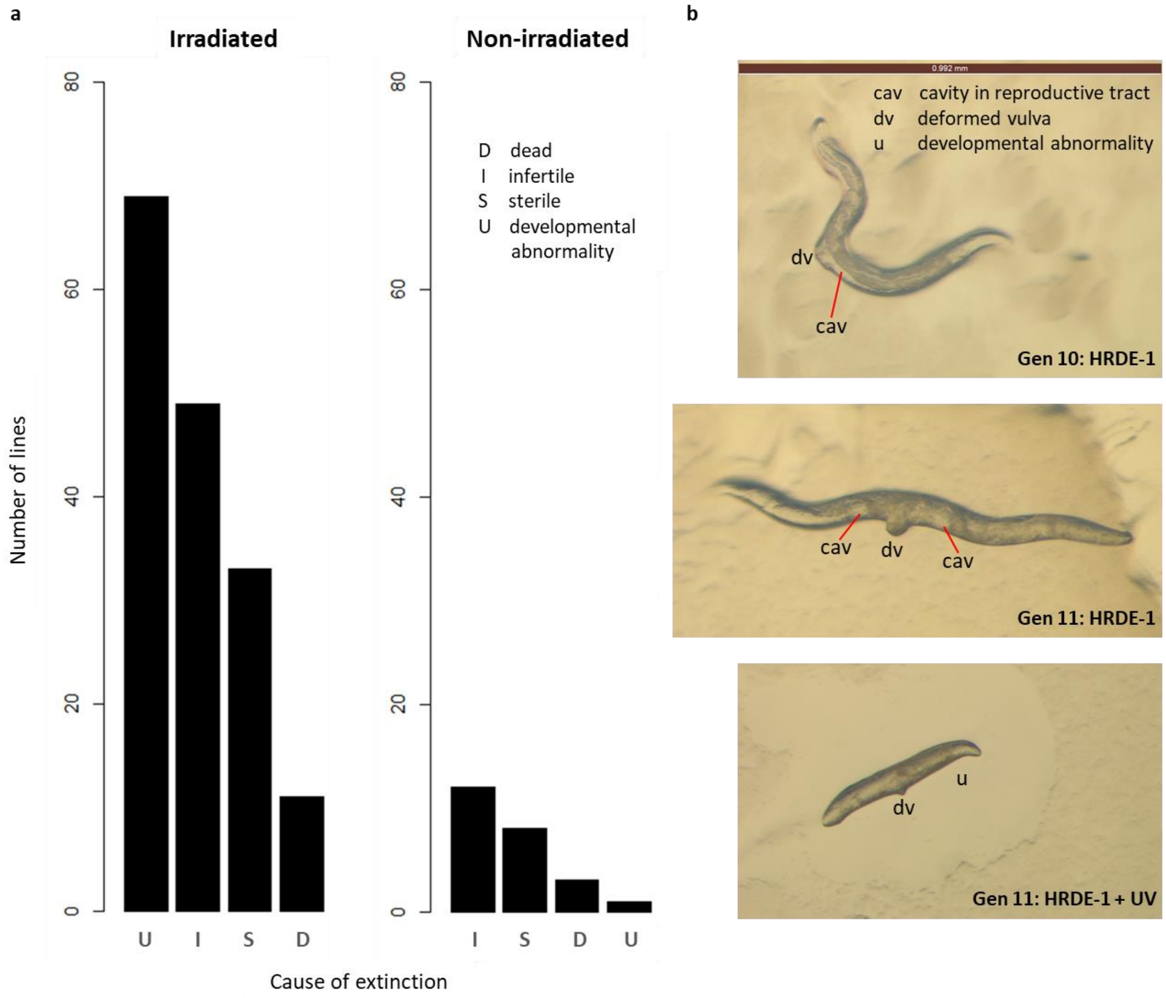


**Supplementary Figure 8: a, Causes of extinction of mutation accumulation (MA) lines in *hrde-1* mutant background. b, Representative images showing impacts of germline damage in *hrde-1* mutant MA lines.** Brown scale bar of 0.992 mm for all images.

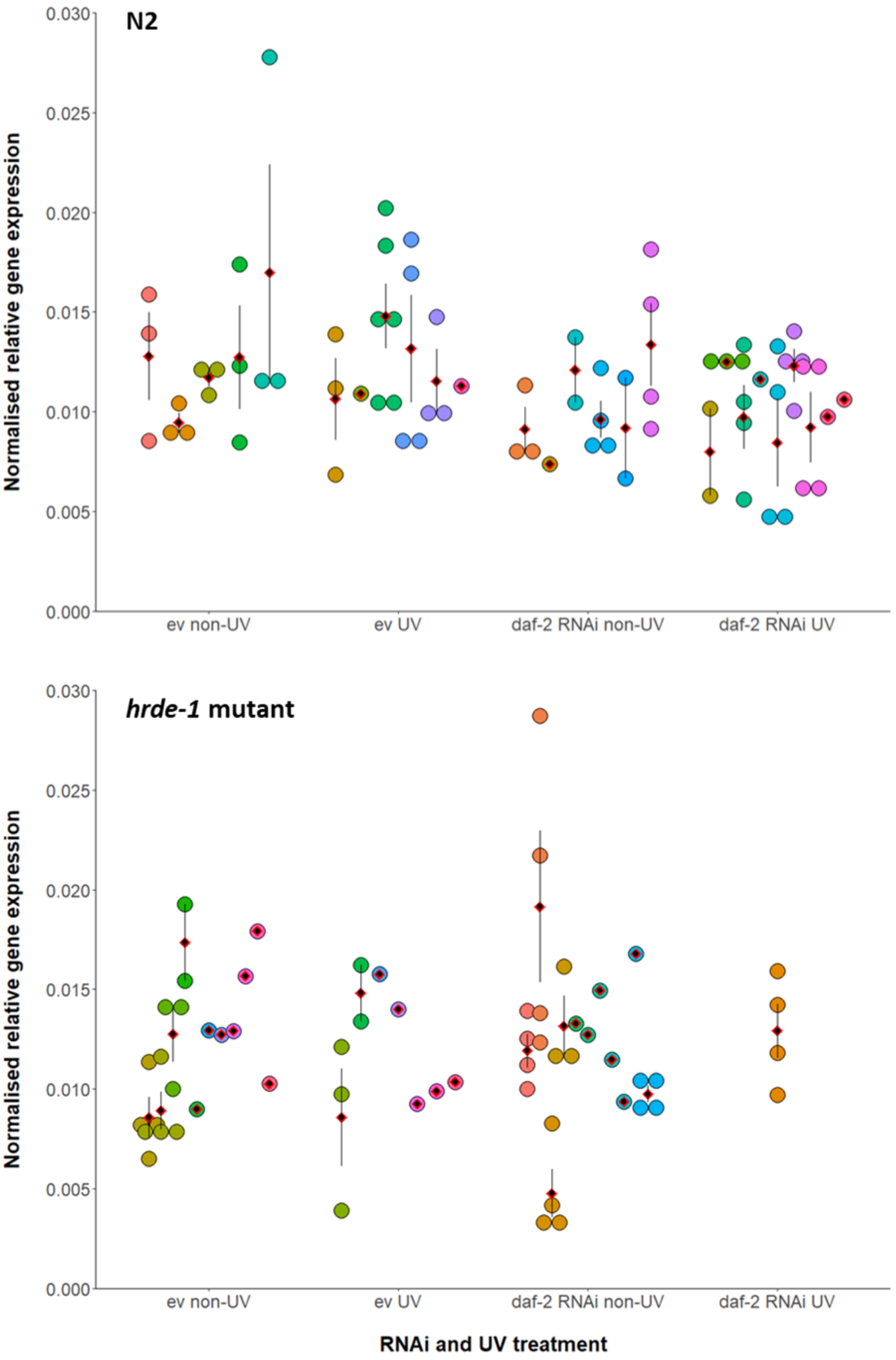

N2
Normalised relative gene expression
0.030
0.025
0.020
0.015
0.010
0.005
0.000
ev non-UV
ev UV
daf-2 RNAi non-UV
daf-2 RNAi UV
hrde-1 mutant
Normalised relative gene expression
0.030
0.025
0.020
0.015
0.010
0.005
0.000
ev non-UV
ev UV
daf-2 RNAi non-UV
daf-2 RNAi UV
RNAi and UV treatment

**Supplementary Figure 9. Normalised *daf-2* expression following *daf-2* RNAi treatment versus untreated empty vector (ev) controls, from mutation accumulation (MA) lines at generation 20.** UV treatment indicates the background of spontaneous (non-UV) or UV-induced (UV) MA from N2 and *hrde-1* mutant genotypes. RNAi was delivered from late-L4 stage and gene expression quantified in Day 2 adults using qRT-PCR. Arithmetic mean of biological replicates per MA line shown as a red diamond with +/- 1 standard error bars. Each colour represents a different MA line for each treatment combination, from which n=1-6 individual worms (separate points) were assayed. Normalised *daf-2* expression ($2^{-\Delta CT}$) was calculated relative to expression of the *actin-3* reference gene[92].

**Supplementary Table 1:** Sample sizes for gene expression assay. Total number of individual worms assayed for which N=1-6 individual worms were measured per MA line, for the indicated number (N) of MA lines per treatment: N2 or *hrde-1* mutant backgrounds, for UV-induced or spontaneous (non-UV) MA, on *daf-2* RNAi or empty vector (ev) control bacteria.

| MA treatment | Total N | N, MA lines |
|---|---|---|
| N2 non-UV ev | 15 | 5 |
| N2 non-UV *daf-2* RNAi | 16 | 5 |
| N2 UV ev | 18 | 6 |
| N2 UV *daf-2* RNAi | 24 | 8 |
| *hrde-1* mutant non-UV ev | 22 | 10 |
| *hrde-1* mutant non-UV *daf-2* RNAi | 25 | 10 |
| *hrde-1* mutant UV ev | 8 | 6 |
| *hrde-1* mutant UV *daf-2* RNAi | 4 | 1 |

**Supplementary Table 2a:** Full output of zero-inflated generalised Poisson model used to analyse age-specific reproduction data from the first (F1), second (F2) and third (F3) generation of offspring produced by parents treated with *daf-2* RNAi or an empty vector control, in wild-type N2 or *hrde-1* mutant genotypes.

| | z | p |
|---|---|---|
| **F1 generation:** | | |
| RNAi x age | 2.376 | **0.0175*** |
| RNAi x genotype | -1.584 | 0.113 |
| genotype x age | 2.627 | **0.00861*** |
| genotype x age$^2$ | -4.257 | **<0.001*** |
| **F2 generation:** | | |
| RNAi | 0.137 | 0.891 |
| RNAi x genotype | 0.613 | 0.540 |
| genotype x age | 4.615 | **<0.001*** |
| genotype x age$^2$ | -5.309 | **<0.001*** |
| **F3 generation:** | | |
| RNAi | 0.656 | 0.512 |
| RNAi x genotype | -0.853 | 0.393 |
| genotype x age | 5.630 | **<0.001*** |
| genotype x age$^2$ | -6.383 | **<0.001*** |

**Supplementary Table 2b:** Full output of generalised linear model with Gaussian error structure used to analyse total reproduction data from the first (F1), second (F2) and third (F3) generation of offspring produced by parents from part **a**.

| | t | df | p |
|---|---|---|---|
| **F1 generation:** | | | |
| RNAi x genotype | -2.888 | 1 | **0.00464*** |
| **F2 generation:** | | | |
| RNAi | 0.286 | 1 | 0.775 |
| genotype | 3.787 | 1 | **<0.001*** |
| RNAi x genotype | 1.309 | 1 | 0.193 |
| **F3 generation:** | | | |
| RNAi | 0.380 | 1 | 0.704 |
| genotype | 3.890 | 1 | **<0.001*** |
| RNAi x genotype | -0.746 | 1 | 0.457 |

**Supplementary Table 3a: Relative gene expression (ΔCt) in N2 MA lines.** The effect of *daf-2* RNAi from the late-L4 stage on the expression of *daf-2* relative to the *actin-3* reference gene (ΔCt from qRT-PCR) at day 2 of adulthood, after 20 generations of MA. RNAi and UV treatment contrasts from linear mixed effect model analysis of variance table.

| Factor | Sum Sq | Mean Sq | F | d.f. | p |
|---|---|---|---|---|---|
| RNAi treatment | 1.238 | 1.238 | 6.168 | 1 | **0.001** |
| UV treatment | 0.014 | 0.014 | 0.071 | 1 | 0.793 |
| RNAi x UV treatment | 0.021 | 0.021 | 0.107 | 1 | 0.747 |

**Supplementary Table 3b: Relative gene expression (ΔCt) in *hrde-1* mutant MA lines.** The effect of *daf-2* RNAi from the late-L4 stage on the expression of *daf-2* relative to the *actin-3* reference gene (ΔCt from qRT-PCR) at day 2 of adulthood. RNAi treatment contrast from linear mixed effect model analysis of variance table. Data for non-irradiated *hrde-1* mutant background, after 20 generations of MA.

| Factor | Sum Sq | Mean Sq | F | d.f. | p |
|---|---|---|---|---|---|
| RNAi treatment | 0.012 | 0.012 | 0.094 | 1 | 0.7618 |

**Supplementary Table 3c. Relative gene expression (ΔCt) in combined analysis of N2 and *hrde-1* mutant MA lines.** RNAi, UV and Genotype treatment contrasts and their interactions from linear mixed effect model analysis of variance table.

| Factor | Sum Sq | Mean Sq | F | d.f. | p |
|---|---|---|---|---|---|
| RNAi treatment | 0.414 | 0.414 | 2.333 | 1 | 0.134 |
| UV treatment | 0.007 | 0.007 | 0.038 | 1 | 0.846 |
| Genotype | 0.032 | 0.032 | 0.183 | 1 | 0.671 |
| Genotype x RNAi | 0.074 | 0.074 | 0.420 | 1 | 0.520 |
| RNAi x UV | 0.0002 | 0.0002 | 0.001 | 1 | 0.972 |

**Supplementary Table 4. Primer sequences**. Forward (fwd) and reverse (rev) sequences are listed in the 5' to 3' direction. Sequences acquired from[89] for *daf-2* and[93] for the *actin-3* reference gene.

| Gene | Primer Sequences |
|---|---|
| *daf-2* | Fwd: GTGGCGTGAGAATGAAGTGAG<br>Rev: GGCTTATCGGCTACAATCGTC |
| *actin-3* | Fwd: CCAAGAGAGGTATCCTTACCCTCAA<br>Rev: AAGCTCATTGTAGAAGGTGTGATGC |

### Supplementary Methods

We conducted five experiments to our test our main hypotheses about the effects of reduced insulin/IGF-1 signalling (IIS) in adulthood, via adult-only *daf-2* RNAi, on the soma and the germline:

1. Inter-generational effects of reduced IIS in parents under UV-induced stress, on parental and offspring fitness and reproduction.

2. Effects of reduced IIS on parental fitness and total reproduction under intermittent fasting.

2. Effects of reduced IIS on 40 generations of spontaneous and UV-induced mutation accumulation, in N2 wild-type and RNAi inheritance deficient (*hrde-1*) mutant backgrounds.

3. Life history and fitness effects of 20 generations of spontaneous and UV-induced mutation accumulation under reduced adulthood IIS, on the surviving MA lines.

4. Transgenerational effects of *daf-2* RNAi on offspring fitness in N2 and *hrde-1* mutant backgrounds.

#### Nematode stocks and culture

All experimental lines were kept at 20°C, 60% relative humidity and in darkness, consistent with standard *C. elegans* rearing protocol[94]. *C. elegans* is a valuable model system due to its short life cycle, ease of genetic manipulation and its normal reproductive state as self-fertilising hermaphrodites. Males were excluded from our experiments and occur at a very low frequency of approximately 0.3% under benign lab conditions and in nature[95,96].

During all experiments, worms were kept on 35mm NGM agar plates (supplemented with 1mM IPTG and 50μg/mL of antibiotic ampicillin, to inhibit the growth of bacteria other than our antibiotic resistant *E. coli*) and seeded with 0.1mL of the e.v. control or *daf-2* RNAi bacteria, 24 to 48 hours before use, for *ad libitum* bacterial growth. Bacterial cultures were prepared prior to the experiments, by growing in LB supplemented with 50μg/mL ampicillin (as[97]).

**Reducing IIS via *daf-2* RNAi feeding in adulthood**

RNAi treatment was applied from the late-L4 stage, immediately prior to the onset of adult sexual maturity and all individuals developed on empty vector (e.v.) control *E. coli* prior to this stage. Normal development, which requires functional *daf-2*, was therefore unaffected[19,31].

**Validation of *daf-2* gene expression under RNAi**

Worms were collected on day two of adulthood, to assay the downregulation of *daf-2* in individuals at peak reproduction. To do this we picked individual worms onto unseeded plates and allowed them to crawl around to remove surface bacteria and separate day two adults from their eggs[89], and to ensure ingested bacteria was excreted[98]. This avoided the possibility of any residual RNAi bacteria carrying through to contaminate the RT-qPCR analysis, as[89].

Individual cleaned worms were transferred into 10 μl of worm lysis buffer (containing 1:100 diluted DNase; both from Ambion Power SYBR Green Cells-to-Ct kit) in the separate domed lids of 0.2 ml PCR tubes, immediately spun down and flash frozen in liquid nitrogen[89]. To crack the tough nematode cuticle and release nucleic acids, we performed 10 freeze-thaw cycles, by transferring the PCR tubes between liquid nitrogen and a 40°C, before homogenising the samples in a thermal mixer (Eppendorf ThermoMixer C) set at 4°C, for 30 minutes at 1800 rpm[89]. We confirmed, using Nanodrop spectrometry (Thermo Scientific) that it was possible to obtain ~30-55 ng of RNA from our single worm samples (as[89,99]).

We reverse transcribed DNase-treated RNA using the Ambion Power SYBR Green Cells-to-Ct kit, following the manufacturer protocol. We included a no reverse transcriptase control (NRTC) per RNAi treatment, for which the RT enzyme was substituted with nuclease-free water. The synthesised cDNA was used undiluted for PCR and qRT-PCR.

To confirm that any contaminating genomic DNA had been removed, we performed a standard PCR with a 10 µl reaction and an annealing temperature of 60°C. We ran 5 µl of the PCR reaction on a 1% agarose gel using ethidium bromide and confirmed both the absence of amplification in the NRTCs verifying the successful removal of any contaminating gDNA, and also the successful amplification of cDNA for primer pairs.

The qRT-PCR was performed on an Applied Biosystems 7500 Real-Time PCR System using the Power SYBR Green Cells-to-Ct kit (Ambion) with the following PCR cycle: 95°C for 10 minutes, followed by 40 cycles of: 95°C for 15 s and 60°C for one minute. The total reaction volume was 20 µl of which 4 µl was cDNA. We used primers specific for the target gene of interest (*daf-2*) and for a reference gene- the housekeeping gene, *actin-3* (T04C12.4), commonly used for *C. elegans*[93]. Primer sequences are listed in Table S4. Primers were designed based on MIQUE guidelines[100] and taken from[89] for *daf-2* and[93] for *actin-3.*

Two qRT-qPCR reactions (technical replicates) were carried out per sample per primer pair, to check for repeatability and RNAi treatments were split evenly across the six plates, to control for any minimal plate effects. We also included two negative template controls (nuclease-free water substituted for cDNA) and one NRTC per primer pair per plate, to test for any contamination.

To calculate relative gene expression, we determined ΔCt as the difference between the qRT-PCR cycle thresholds (Ct values) of the target gene of interest (*daf-2*) and the reference gene, for each sample. The arithmetic mean of the Ct values for the two technical replicates per gene, per sample was used in ΔCt calculations. Statistical analyses were performed on ΔCt[89], using a linear mixed effects model with Gaussian error structure (lme4 package), to determine the effect of RNAi treatment (*daf-2* RNAi versus empty vector controls), UV treatment and genotype (N2 or *hrde-1* mutant) and their interaction on relative gene expression. MA line was fitted as a random effect. The Shapiro-Wilk's normality test confirmed that ΔCt values satisfied the normality assumption of the linear models ($W=0.983$, $p=0.428$).

The coefficient of variation (CV, %) in ΔCt between biological replicates for each MA line, and in Ct values between technical replicates per gene, per sample, was calculated as the standard deviation divided by the mean for each comparison[89,101], to determine biological variation in relative gene expression between individual worms and repeatability of the qPCR results, respectively.

To quantify fold change in gene expression ($2^{-\Delta\Delta CT}$), we calculated the difference in the relative levels of mRNA for the target gene of interest compared to the reference gene (ΔCt) between untreated controls and RNAi treated samples, using mean ΔCt across all biological replicates per RNAi treatment[92].

**Ultraviolet wavelength C (UV-C) irradiation**

We calibrated the UV-C lamp of the Thermo Scientific Heraguard ECO Safety Cabinet to determine the exposure time required to deliver an irradiation dose in the range of previous work for *C. elegans*[45-48]. First, we measured the UV intensity (also known as irradiance or fluence rate) of UV-C radiation emitted at a distance of 66cm from the centre of the lamp to the surface on which our nematodes on plates were placed for irradiation, as 2.3W/$m^2$, using a Macam spectroradiometer and verified with a Sigma Laser Power Energy Meter (Coherent). As the radiation dose is the product of intensity and exposure time (as approved by the International UV Association), we calculated a required exposure time of 20 seconds.

The UV lamp was allowed to warm up and stabilise in temperature for 10 minutes, before irradiation of worms, as UV intensity increases with initial increases in temperature of the UV lamp, but plateaus after at least five minutes[102].

Worms were irradiated on their *E. coli* seeded plates, without lids (plastic would otherwise absorb UV irradiation) and positioned equidistant from the centre of the lamp, in batches of maximum 30 plates, to ensure radiation dose did not differ significantly between plates.

UV irradiation was timed at exactly 24 hours (+/- 30 minutes) after the RNAi treatment began at the onset of adulthood and prior to peak reproduction, to allow time for reduced IIS in individuals on *daf-2* RNAi[44,103]. This timing allowed us to induce germline mutagenesis, as adult somatic tissue in *C. elegans* is post-mitotic and very resistant to irradiation, whereas germline tissue (eggs, developing oocytes and germline stem cells) is still actively dividing (undergoing meiosis and mitosis) and so is more sensitive to UV irradiation[36]. As spermatogenesis is completed during the late-L4 stage[104], it would not have been directly affected by irradiation[105].

Nematodes were transferred to new seeding immediately after UV irradiation, in case the seeding or plate was affected by UV irradiation, and worms were placed just outside the seeding on the new plate, to minimise contamination with mutated bacteria (as[106,107]). UV-irradiated worms were allowed 8 hours (+/- 30 minutes) recovery, to provide sufficient time for expulsion of irradiated embryos, before starting egg laying[47,107]. Eggs laid were therefore most likely irradiated either as oocytes, or as germline stem cells[104,107].

**Inter-generational effects of UV irradiation and reduced IIS in parents on parental and offspring fitness and reproduction**

Wild-type N2 parents were either UV-irradiated or not as Day 2 adults and maintained on *daf-2* RNAi or an empty vector control for the whole of adulthood, in fully factorial design. Offspring were all non-irradiated and maintained on empty vector throughout life. We assayed the daily offspring production from unmated, singly-held hermaphrodite parents, for their entire reproductive period (first six days of adulthood) and also from the first generation of their offspring, by daily transfers to fresh plates. To assess parental survival, daily mortality checks were made, with death being defined as no observed movement after gentle prodding. Worms were grouped as ten worms per plate after the six-day reproductive period, for logistical reasons and these groups were maintained as independent non-mixing units, for daily transfers across lifetime so that plate identity could be included as a random effect in analysis. One non-irradiated worm on *daf-2* RNAi did not produce any offspring and was omitted from the analysis.

**Effects of reduced IIS on parental fitness and total reproduction under intermittent fasting**

N2 wild type adults were intermittently fasted (IF) by transferring to unseeded IPTG plates without peptone (to avoid bacterial growth) during the 9h starvation periods. Individuals from both ad libitum (AL) and IF treatments were fed empty vector AL during development and then half of each AL and IF treatment were switched to *daf-2* RNAi from late-L4 onwards whilst the other half were maintained on ev (n=100 individuals per treatment, across two time-staggered blacks of n=50 per treatment). IF individuals were fed AL outside of 9h starvations. Adults were assayed for daily reproduction to calculate lambda fitness and total lifetime reproduction.

**Mutation accumulation (MA) design**

Each generation, for all treatments, we allowed mid-day 2 to mid-day 3 adults to lay eggs onto empty vector plates, from which we picked a single late-L4 larva per line to form the next generation of MA. The late-L4 stage is easily identifiable in *C. elegans* (vulva cells are visible in the vulva lumen and prior to vulval protrusion at sexual maturity), which allowed repeatable and consistent age-controlled set-up of each generation. The duration of the egg lay period was optimised, to account for differences in developmental timing between treatments. As such, even though the development time was longer for eggs laid by parents from UV-irradiated MA lines and for all treatments with increasing MA generation, we limited selection on development time unevenly between treatments, as the next MA generation was set-up at an identical late-L4 developmental stage for all treatments. Age of parental egg lay was alternated between mid-Day 2 and mid-Day 3 every second generation, to limit selection on parental age at reproduction and so the offspring forming each new MA generation came from parents during their period of peak reproduction, when most individuals were reproducing.

We recorded the generation at which extinction occurred and the cause of extinction (death, failure to produce viable eggs or failure to reach sexually mature adult). Individuals that were lost, underwent matricide (internal hatching that killed the parent) prior to the Day 2 egg lay, died from the expulsion of internal tissue, became infected or desiccated on the plate wall were censored. We photographed a sample of Day 1 adult worms with visible developmental or reproductive abnormalities under light microscopy (camera specifications in egg size measurement section below), including worms with stunted growth, apparent aberrant external growth (tumour), deformed vulva, or abnormal cavities in the reproductive tract in place of oocytes or embryos.

**Effect of 20 generations of spontaneous and UV-induced mutation accumulation on fitness and life history of surviving MA lines**

Prior to the life history assay, individuals taken from MA lines at generation 20 were reared for two subsequent generations under common garden conditions (no irradiation, on empty vector control), to reduce direct parental effects from exposure to RNAi and UV, and hence look for genetic differences between MA treatments. The life history assay was conducted in the common garden environment.

**Age-specific reproduction, fitness and total lifetime reproduction assays**

The first 4 days of adult reproduction were used to calculate fitness and to analyse age-specific reproduction, as day 5 and 6 reproduction was zero for almost all individuals. Parental fitness and age-specific reproduction were analysed for day 2 to 4 inclusive, for the first intergenerational experiment, as UV-irradiation was administered at the start of day 2 of adulthood.

**Egg size measurement**

Only eggs laid at the normal gastrula (approximately 30-cell) stage of development were photographed, as egg shape and size can vary after 9 hours of ex-utero development. Egg size (area in $mm^2$ enclosed within a free-drawn ellipse around the egg perimeter) was calculated from photos using *Image J Fiji program* v.1.51[108], whilst blind to treatment identity.

To minimise experimenter error, each egg size measurement was taken twice and an average taken, and the same experimenter measured all eggs. Egg measurement was completed across three measurement days, using identical ImageJ settings and scale calibration. Treatments and blocks were stratified across measurement days and randomly dispersed within each day, to avoid bias. Egg area measurements were strongly repeatable between replicate measurements on the same egg (Spearman's rank correlation coefficient, rho= 0.899), there was no temporal autocorrelation between measurement days (Kruskal-Wallis rank sum test, $\chi^2$=299, df=314, p=0.721) and only weak temporal autocorrelation on the first measurement day (Spearman's rank, rho=0.398).

**Male production**

We determined the percentage of eggs that developed into adult males (n= 75 eggs assayed/treatment), one egg per individual Day 2 worm, as above, to determine if the stress induced by irradiation or mutation accumulation would increase male production above standard N2 wild-type levels, as seen previously for heat stress and starvation in *C. elegans*[76,77].

**Adult heat shock resistance**

To test the stress resistance of post-reproductive (Day 7) adults, we assayed survival following acute heat shock of 1 hour and 45 minutes at 37°C (following[83]), in the same individuals used for the Generation 20 age-specific reproduction assay. At 3 and 24 hours post-heat shock we recorded any individuals that had died and categorised locomotion as normal movement (crawling spontaneously or when gently prodded), uncoordinated movement of head and tail with no forward or backward trajectory, or just head movements; following[109,110]. Previous work confirmed that from 12 hours after heat shock is a reliable time point to score long-term survival in *C. elegans* adults[83]. All worms were heat-shocked simultaneously, grouped by treatment, five worms per plate and the positions of plates in the oven were stratified by treatment, to control for any positional differences in heat exposure.

**Statistical analyses**

The 'dabest' Cummings estimation plots display all datapoints from each treatment as a swarm-plot, with the mean ± standard deviation of treatment as a gapped line adjacent to the points. The mean difference (effect size) and its 95% confidence interval (CI) is estimated for pairwise comparisons of treatments via non-parametric bootstrap resampling (n=5000) and displayed below the main plots, for visualisation. When the 95% CI vertical bar does not cross the x-axis (mean difference equal to zero), there is a significant difference between the pairwise treatment comparison. Individual fitness (lambda) incorporates age-specific reproduction, with emphasis on early reproduction and fast development. It is therefore an appropriate measure when timing is important for fitness[55]. Individual fitness was calculated from the life table of age-specific reproduction, by solving the Euler-Lotka equation using the lambda function in the popbio package in R (as[20,90]). The life table was constructed using a common development time of two days without reproduction. Worms matured and reproduction began during day three (first day of adulthood) when fast-developing worms would have matured earlier and have had more reproduction on this day. Since lambda is a rate-sensitive measure, fast development, resulting in increased early life reproduction, would translate into higher values of lambda.

Fitness and total reproduction data from the intermittent fasting assay were analysed using a generalised linear mixed effects model (GLM) with Gaussian error structure ('lmer' function in 'lme4' package), fitting diet regime (either intermittent fasting, IF or ad libitum feeding, AL), RNAi treatment (ev or *daf-2* RNAi) and their interaction as fixed effects and block as a random effect. Individuals which were lost due to walling were excluded from the analysis (exclusions per treatment: AL ev n=1, AL *daf-2* RNAi n=1, IF ev n=26 and IF *daf-2* RNAi n=27).
